# Architecture and evolution of subtelomeres in the unicellular green alga *Chlamydomonas reinhardtii*

**DOI:** 10.1101/2021.01.29.428817

**Authors:** Frédéric Chaux-Jukic, Samuel O’Donnell, Rory J. Craig, Stephan Eberhard, Olivier Vallon, Zhou Xu

## Abstract

In most eukaryotes, subtelomeres are dynamic genomic regions populated by multi-copy sequences of different origins, which can promote segmental duplications and chromosomal rearrangements. However, their repetitive nature has complicated the efforts to sequence them, analyze their structure and infer how they evolved. Here, we use recent and forthcoming genome assemblies of *Chlamydomonas reinhardtii* based on long-read sequencing to comprehensively describe the subtelomere architecture of the 17 chromosomes of this model unicellular green alga. We identify three main repeated elements present at subtelomeres, which we call *Sultan*, *Subtile* and *Suber*, alongside three chromosome extremities with ribosomal DNA as the only identified component of their subtelomeres. The most common architecture, present in 27 out of 34 subtelomeres, is an array of 1 to 46 tandem copies of *Sultan* elements adjacent to the telomere and followed by a transcribed centromere-proximal *Spacer* sequence, a G-rich microsatellite and a region rich in transposable elements. Sequence similarity analyses suggest that *Sultan* elements underwent segmental duplications within each subtelomere and rearranged between subtelomeres at a much lower frequency. Comparison of genomic sequences of three laboratory strains and a wild isolate of *C. reinhardtii* shows that the overall subtelomeric architecture was already present in their last common ancestor, although subtelomeric rearrangements are on-going at the species level. Analysis of other green algae reveals the presence of species-specific repeated elements, highly conserved across subtelomeres and unrelated to the *Sultan* element, but with a subtelomere structure similar to *C. reinhardtii*. Overall, our work uncovers the complexity and evolution of subtelomere architecture in green algae.

## Introduction

The extremities of linear chromosomes in eukaryotes are essential to maintain stable genomes (Jain and Cooper 2010). At their very end, repeated sequences called telomeres recruit specific factors that collectively prevent detection of the extremities as double-strand breaks and avoid deleterious effects caused by repair attempts by the cell (Wellinger and Zakian 2012; de Lange 2018). Telomeres also counteract the end replication problem, which would otherwise lead to replicative senescence and cell death. In most organisms, this is achieved by recruiting the reverse-transcriptase telomerase, which processively adds *de novo* telomere sequences. Instead of telomerase, some species of the Diptera order use other maintenance mechanisms, such as retrotransposons in *Drosophila melanogaster* or recombination-dependent mechanisms in *Chironomus* or *Anopheles* (Cohn and Edstrom 1992; Roth et al. 1997; Pardue and DeBaryshe 2011). Homology-directed recombination can also be used to maintain telomeres in a number of cancer cells and in experimental models where telomerase is inactivated (Cesare and Reddel 2010). Next to the telomere, the subtelomere is commonly a gene-poor region comprising repeated elements, such as transposable elements (TEs), satellite sequences, or paralogous genes, which are often shared between different subtelomeres (Corcoran et al. 1988; Louis 1995; Kim et al. 1998; Fabre et al. 2005; Brown et al. 2010; Richard et al. 2013; Chen et al. 2018). In some organisms, these families of non-essential paralogous genes are involved in growth and response to specific environments, and subtelomeres have been proposed to be a nursery for new genes (Wickstead et al. 2003; Fabre et al. 2005; Brown et al. 2010; Chen et al. 2018). Although subtelomeres are mostly heterochromatic (Gottschling et al. 1990; Baur et al. 2001; Pedram et al. 2006; Jain et al. 2010; Vrbsky et al. 2010), specific transcripts have been detected in these regions, including the telomeric repeat-containing RNA (TERRA), which plays multiple roles in telomere biology (Azzalin et al. 2007; Azzalin and Lingner 2015). Importantly, subtelomeres can regulate telomere length, telomere-associated chromatin, replicative senescence and help maintain telomere and genome integrity (Gottschling et al. 1990; Craven and Petes 1999; Fabre et al. 2005; Arneric and Lingner 2007; Azzalin et al. 2007; Schoeftner and Blasco 2008; Tashiro et al. 2017; Jolivet et al. 2019).

Subtelomeres are rapidly evolving regions and can vary greatly in structure and composition between closely related species and even individuals of the same species (Horowitz and Haber 1984; Louis and Haber 1992; Louis et al. 1994; Anderson et al. 2008; Rudd et al. 2009; Yue et al. 2017; Kim et al. 2019; Young et al. 2020). Several mechanisms have been shown or proposed to explain subtelomeric variations. The repetitive nature of the region promotes homologous recombination (HR), unequal sister chromatid exchange (SCE), break-induced replication (BIR) and replication slippage (Horowitz and Haber 1984; Corcoran et al. 1988; Louis and Haber 1990; Linardopoulou et al. 2005; Kuo et al. 2006; Rudd et al. 2007; Wang et al. 2010; Chen et al. 2018; Kim et al. 2019). Transposition also contributes to subtelomere variations (Kim et al. 1998; Kuo et al. 2006; Rudd et al. 2009; Chen et al. 2018). All of these mechanisms, along with others such as non-homologous end-joining (NHEJ)-mediated translocations and fusions, can lead to segmental duplications and amplification of repeated elements (Linardopoulou et al. 2005; Kuo et al. 2006; Wang et al. 2010; Chen et al. 2018). Consistently, mutation rates and chromosomal rearrangements are elevated at chromosome ends, even more so in the absence of telomerase (Horowitz and Haber 1984; Hackett et al. 2001; Siroky et al. 2003; Londono-Vallejo et al. 2004; Anderson et al. 2008; Coutelier et al. 2018).

Subtelomeres are therefore of critical importance for both genome stability and evolution. But because of their intrinsically complex and repetitive nature, telomeres and subtelomeres are often misassembled or altogether absent in reference genomes of most species. For example, the human reference genome still lacks a comprehensive and accurate representation of its subtelomeres, although recent advances improved the assembly (Stong et al. 2014; Logsdon et al. 2020; Miga et al. 2020; Young et al. 2020). With the advent of long read sequencing technologies (Li et al. 2017; Yue et al. 2017; Kim et al. 2019), we can look forward to better assemblies and descriptions of subtelomeres, enabling the mechanisms underlying their structural variations and evolution to be inferred for a diverse range of organisms.

We recently characterized telomere structure and telomerase mutants in the unicellular green alga *Chlamydomonas reinhardtii* (Eberhard et al. 2019), a major model for photosynthesis and cilia research. The discovery of blunt ends at a subset of telomeres and a wide range of telomere length distributions in different laboratory strains and natural isolates prompted us to further explore how chromosome ends have evolved and are structured. Here, we provide a comprehensive description of the architecture of the subtelomeres in *C. reinhardtii* and a comparative analysis with other green algae. An early study evidenced a high level of similarity in the sequences adjacent to a few cloned *C. reinhardtii* telomeres (Petracek et al. 1990). Subtelomere architecture has also been partially outlined in a limited number of plant species, including *Arabidopsis thaliana*, *Silene latifolia* and *Phaseolus vulgaris* (Kotani et al. 1999; Sykorova et al. 2003; Kuo et al. 2006; Wang et al. 2010; Richard et al. 2013; Chen et al. 2018), and the green alga *Coccomyxa subellipsoidea* (Blanc et al. 2012). To probe the structure of these repetitive regions, we recently generated a contiguous *de novo* assembly from published Oxford Nanopore Technologies long reads (Liu et al. 2019; O'Donnell et al. 2020), which we analyze alongside soon-to-be-released PacBio-generated assemblies (Craig et al., *in prep*). We show that most *C. reinhardtii* subtelomeres are composed, reading from the telomere toward the centromere, of an array of repeated elements that we call *Sultan* (for *SUbtelomeric Long TANdem repeats*), a *Spacer* sequence, a variable number of G-rich microsatellite sequences and various types of TEs. Sequence homology analysis of the *Sultan* elements suggests that they mostly propagated within a subtelomere through segmental duplications and less frequently between different subtelomeres. Subtelomeres in other green algae also contain specific repeated sequences, unrelated to the *Sultan* element, suggesting a common structure that has possibly evolved independently and is important for subtelomere function.

## Results

### *Chlamydomonas* subtelomeres comprise arrays of specific tandemly repeated sequences

Because the publicly available *C. reinhardtii* reference genome version 5 (v5; https://phytozome-next.jgi.doe.gov/) at the time of this work was incompletely assembled near the chromosome extremities, we took advantage of our recent release of a *de novo* genome assembly (O'Donnell et al. 2020) based on long-read sequencing data of strain CC-1690 (Liu et al. 2019), a commonly used laboratory strain also known as 21gr. Briefly, the raw Oxford Nanopore Technologies (hereafter referred to as “Nanopore”) electrical signal was base-called and subsets of the longest reads (*N50* ⋍ 55 kb) were assembled independently using various protocols, after which assemblies were combined to create a 21-contig genome assembly, readily scaffolded onto the 17 chromosomes. The chromosome arms, and therefore their termini, are labelled “left” and “right” (or _L and _R) based on the orientation used in the reference genome and the sequences and features are generally presented reading from the telomere towards the centromere.

Arrays of the 8-bp telomeric repeat motif (5’-CCCTAAAA-3’/5’-TTTTAGGG-3’) previously described in *C. reinhardtii* (Petracek et al. 1990; Fulneckova et al. 2012; Eberhard et al. 2019) were found at the extremity of 31 out of the 34 chromosome ends (Fig. 1, black segments). In the genome assembly, telomeric repeats had a median length of 311 ± 125 bp (median ± SD), at the shorter end of the 300-700 bp range observed previously by terminal restriction fragment analysis (Eberhard et al. 2019).

**Figure 1.**
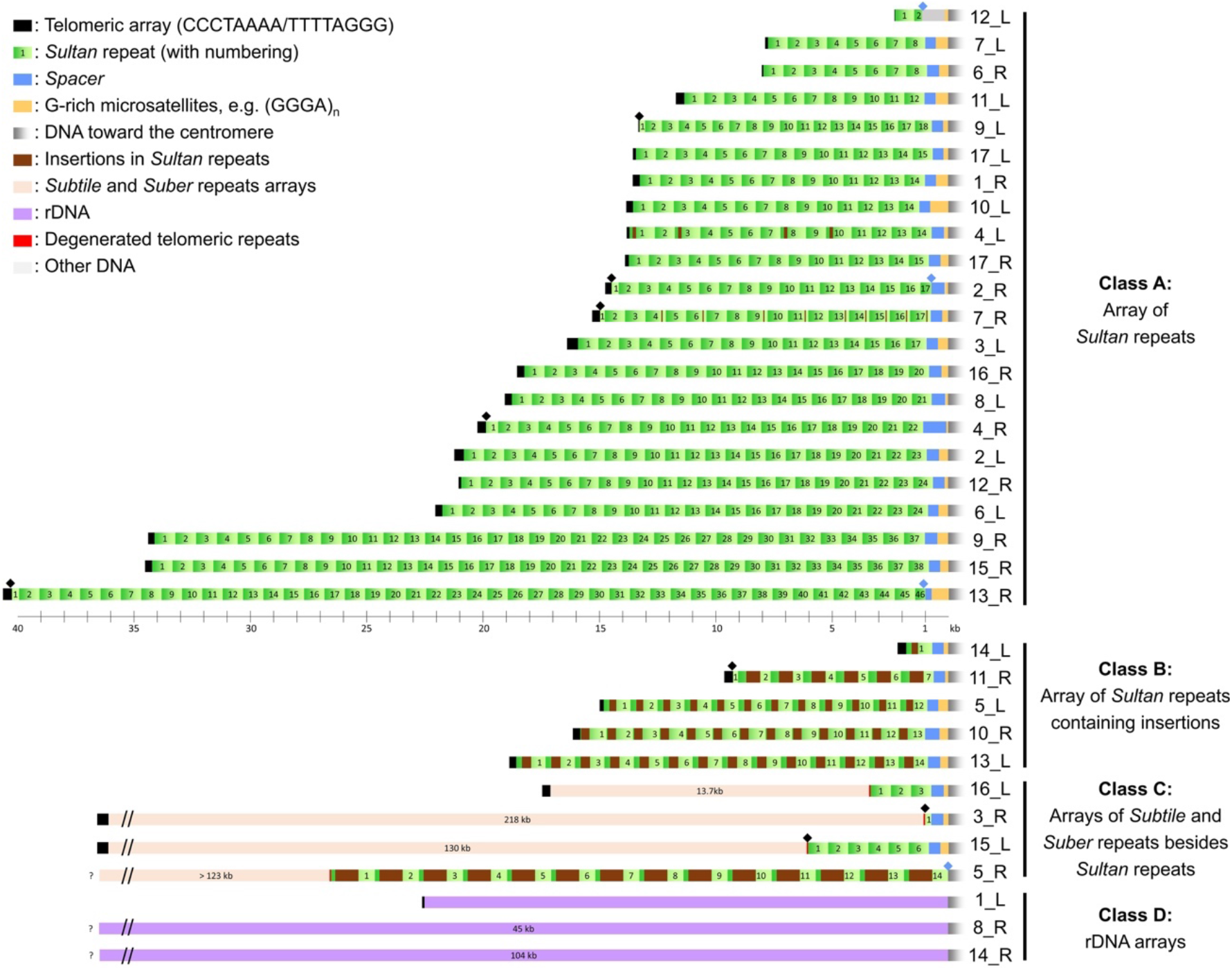
Architecture of subtelomeres in *C. reinhardtii* strain CC-1690. Left and right ends (_L and _R, respectively) of CC-1690 chromosomes are depicted with telomeres on the left-hand side, sorted by class and number of subtelomeric elements, which are displayed as boxes drawn at scale. The most common architecture, class A subtelomeres, comprises a telomere tract (black), a tandem array of *Sultan* repeats (green; numbering starts on the telomere side), a *Spacer* sequence (blue) and a G-rich microsatellite (yellow). Distinct large DNA insertions (brown) found in the *Sultan* repeats define the class B subtelomeres. Other repeats (pink) are found upstream of the *Sultan* array in class C subtelomeres (see Fig. 4). Arrays of ribosomal DNA (purple) compose class D subtelomeres (Supplemental Figure S1). The display of the longest class C and D subtelomeres is not at scale and interrupted by “//”, while “?” marks elusive molecule ends due to assembly collapse. Diamonds denote junctions with telomere or *Spacer* interrupting a *Sultan* element (see Supplemental Table ST1).

Alignments systematically revealed extensive homology between subtelomeres, usually covering several kilobases in the form of long repeated arrays. To identify the repeated elements in subtelomeres, we scanned the last 30 kb of each chromosome end using XSTREAM (Newman and Cooper 2007) and Tandem Repeat Finder (TRF) (Benson 1999). Figure 1 shows a map of the subtelomeres of strain CC-1690, depicting their repetitive architecture and shared elements. The most widespread arrays were composed of a ~850 bp element, repeated in direct orientation without interspersed sequence and absent from the rest of the genome, which we thus called *Sultan* for *SUbtelomeric Long TANdem* repeat (Fig. 1, green boxes).

We categorized all subtelomeres into 4 classes. The 27 subtelomeres containing *Sultan* arrays adjacent to the telomeres belonged either to class A, when the *Sultan* elements overall closely matched the most common ~850 bp sequence, or class B, when the *Sultan* elements carried large insertions (Supplemental File F1). In the 4 class C subtelomeres, *Sultan* arrays are separated from the telomeres by large arrays of other repetitive sequences (Fig. 1, pale pink boxes). Finally, class D extremities 1_L, 8_R and 14_R contained rDNA as the only subtelomeric element (Fig. 1, purple segments; Supplemental Fig. S1). In 1_L, only one partial and one complete rDNA copy (which was disrupted by a retrotransposon) were present, which were capped by a telomere. In contrast, 8_R and 14_R appeared to be full arrays, which have been estimated to carry 250-400 copies (>2000 kb) collectively (Howell 1972; Marco and Rochaix 1980), hence much longer than average Nanopore reads, preventing genome assembly from reaching the actual end of the chromosomes.

The number of *Sultan* copies in an array was highly variable, with an overall median of 14 repeats, leading to vastly distinct array lengths (Supplemental Table ST1). Importantly, to verify that the number of *Sultan* repeats was correctly assessed in class A and B subtelomeres, we manually verified the colinearity between individual long reads from the raw unassembled dataset (Supplemental Fig. S2).

In 29 out of the 31 *Sultan*-containing subtelomeres, we found a non-repetitive sequence adjacent to the most centromere-proximal *Sultan* that we called “*Spacer*” (Fig. 1, blue boxes), since it seemed to connect the *Sultan* array to a (GGGA)_n_ microsatellite (Fig. 1, yellow boxes). Most *Spacer* sequences were 450-550 bp long and on the *Sultan* repeat side, the first dozen nucleotides were highly conserved across subtelomeres (Supplemental File F1, TGGTG**A**G**A**GCAAAC found in 24 subtelomeres and TGGTG**C**G**G**GCAAACATTT found in 4, the two least conserved nucleotides are in bold). Three *Spacers* were different: the one in subtelomere 12_L lost homology to the others after the first 40 nt, 13_R was truncated on the *Sultan* repeat side and 10_R displayed a 140-bp insertion just downstream of the highly conserved start described above. In the deep transcriptome data published by (Strenkert et al. 2019), we found 100%-matching reads in nearly all *Spacer* sequences (Supplemental Fig. S3A and B), with the exception of the 5’ truncated 13_R *Spacer*. Using Iso-Seq data (Gallaher et al. 2021) we observed full-length polyadenylated transcripts originating from the *Spacer* towards the centromere at 14 subtelomeres (Supplemental Fig. S4). Furthermore, we observed peaks in H3K4me3 ChIP-seq coverage (Gallaher et al. 2021) at the *Spacers*, which are highly indicative of transcription start sites and active promoters in *C. reinhardtii* (Ngan et al. 2015). These transcripts were generally characterized by a conserved 5’ splice site (G^GTAG), with the (GGGA)_n_ repeat positioned at the beginning of the first, usually extremely long, intron. Sequence similarity was limited to the first exon. In several cases the *Spacer* appeared to act as an alternative promoter to an independent downstream gene (Supplemental Fig. S4C), and the transcripts originating from the *Spacer* only contained very short ORFs. Interestingly, the expression of *Spacer* sequences peaked at dusk in synchronous diurnal cultures (Supplemental Fig. S3C), correlating with transcription of genes associated with DNA replication (Strenkert et al. 2019).

Since it is shared by 27 out of 34 chromosome extremities, we propose that the canonical architecture of a subtelomere in *C. reinhardtii* is, from telomeres inward, an array of *Sultan* repeats, a *Spacer* sequence and a G-rich microsatellite array.

### *Sultan* element organization and dynamics

To further examine subtelomere architecture, we compared the sequences of all 483 *Sultan* elements pair-wise (Fig. 2A; Supplemental File F1). We found that *Sultan* repeats were systematically more conserved within a given subtelomere than between them, although some subtelomeres, such as 2_L, 2_R, 8_L and 12_L, shared highly similar *Sultan* elements (Fig. 2A).

**Figure 2.**
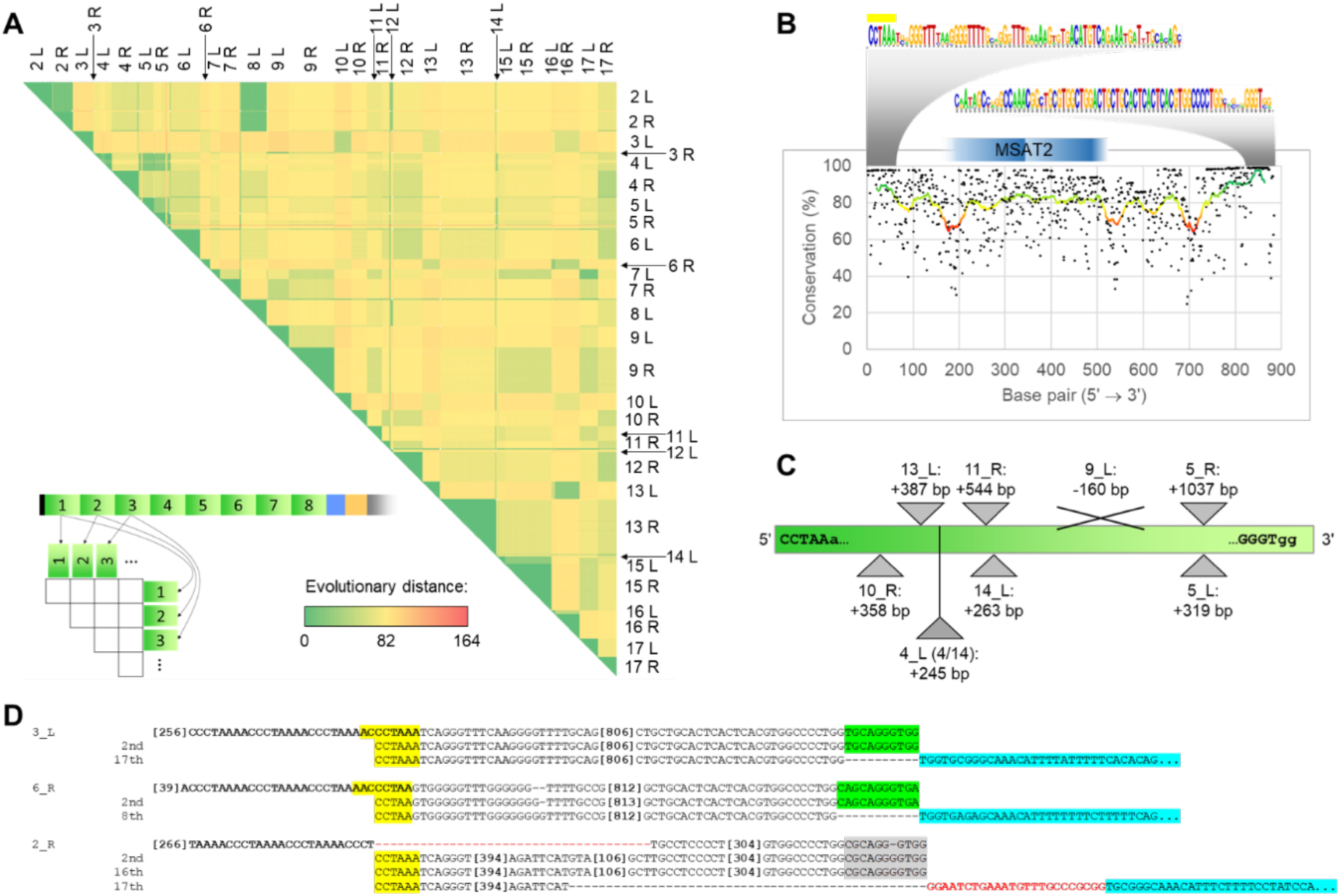
The *Sultan* element. All individual *Sultan* sequences were aligned using MAFFT (Supplemental File F1). *(A)* Pairwise distance heatmap of 483 individual *Sultan* copies. Color scale of the distances, with Jukes-Cantor correction for multiple substitutions, ranges from 0 (green) to 164 (red). Lower left: scheme depicting the numbering of *Sultan* elements within a subtelomere. *(B)* Conservation of nucleotides (black dots) in the consensus sequence. In addition, the moving 40 bp average is plotted as a line (red to green gradient). The sequence logos are shown for the most conserved regions. The telomere-like sequence at start is highlighted in yellow. Similarity with satellite *MSAT2_CR* is indicated in dark blue. *(C)* Location of the largest insertions (triangle) and deletion (X) in *Sultan* repeats of class B subtelomeres (Supplemental Table ST2). *(D)* Alignment of the first and last nucleotides of representative *Sultan* arrays showing phased (3_L and 6_R) and unphased transitions (2_R) to telomere repeats. 5’ telomere-like sequences are shown in yellow, a 10-12-bp sequence in the 3’ region absent in the *Sultan* closest to the *Spacer* is highlighted in green and the 5’ region of *Spacer* in blue. A 22-bp insertion in the transition from *Sultan* to *Spacer* in subtelomere 2_R is shown in red.

*Sultan* elements contained a telomere-like sequence (CCTAAA, CCTAA or CTAAA) on their left border (Fig. 2B). Interestingly, this sequence served as a seamless transition into the telomeric tracts (CCCTAAAA)_n_ on most subtelomeres (Fig. 2D), suggesting that this sequence might act as a seed for telomere elongation. Exceptions are shown by black diamonds in Fig. 1 and exemplified by 2_R in Fig. 2D, where the telomere-proximal *Sultan* lacked > 500 bp as compared to the following repeats. The 3’ side of *Sultan* was also well conserved between chromosomes (Fig. 2B). The last 10-12 nucleotides were truncated in the most centromere-proximal *Sultan* repeat (Fig. 2D), except in rare cases where an insertion or deletion modified the transition from the *Sultan* array to the *Spacer* (Fig. 1, blue diamonds; Fig. 2D, subtelomere 2_R).

We found that the *Sultan* element was poor in GC (average of 53% while genome-wide average is 64%) and their large arrays formed significant regions with lower GC content at the genome-wide level (Supplemental Fig. S5). The central part of *Sultan* sequences was less conserved but showed similarity to the minisatellite *MSAT2_CR* (Fig. 2B), composed of a 184-bp monomer (https://www.girinst.org/2005/vol5/issue3/MSAT-2_CR.html). *MSAT2_CR* was not restricted to subtelomeres and was present in two arrays >10 kb located immediately upstream of the putatively centromeric *Zepp*-like repeats of chromosomes 11 and 13 (Craig et al. 2020). The *Sultan* repeat itself is not a TE: it was detected neither in a previous large-scale survey of TEs, including the terminal repeat in miniature (TRIM) retrotransposons that are shorter than 1000 bp and form long arrays (Gao et al. 2016), nor in a recent annotation of *Chlamydomonas* TEs (Craig et al. 2020), nor in a search against Pfam databases.

In class B subtelomeres, *Sultan* repeats were longer than in class A, due to the presence of large insertions homologous to various TEs (Fig. 2C and Supplemental Table ST2). On a given class B subtelomere, all *Sultan* repeats shared the same inserted element with only minor variations in sequence. The inserted elements were different for each class B subtelomere.

To obtain insights into their propagation, we analyzed the similarity between *Sultan* elements within a subtelomere. On most subtelomeres, individual *Sultan* repeats contain very few variations as compared to the local consensus sequences (> 99.5% identity). Because single-nucleotide variants (SNVs) might result from sequencing and assembly errors, we only used INDELs found in at least two repeats to infer *Sultan* similarity within a given array. The class A subtelomeres 4_L and 7_R harboured an insertion of 245 bp and a duplication of 12 bp, respectively, in a subset of their *Sultan* repeats (Fig. 3). Since in these examples the modified *Sultan* repeats were not contiguous and an identical pattern of modified and standard *Sultans* was found at least twice, we inferred that duplication of *Sultan* elements could involve multiple copies in a single event (Fig. 3B and D, dotted brackets).

**Figure 3.**
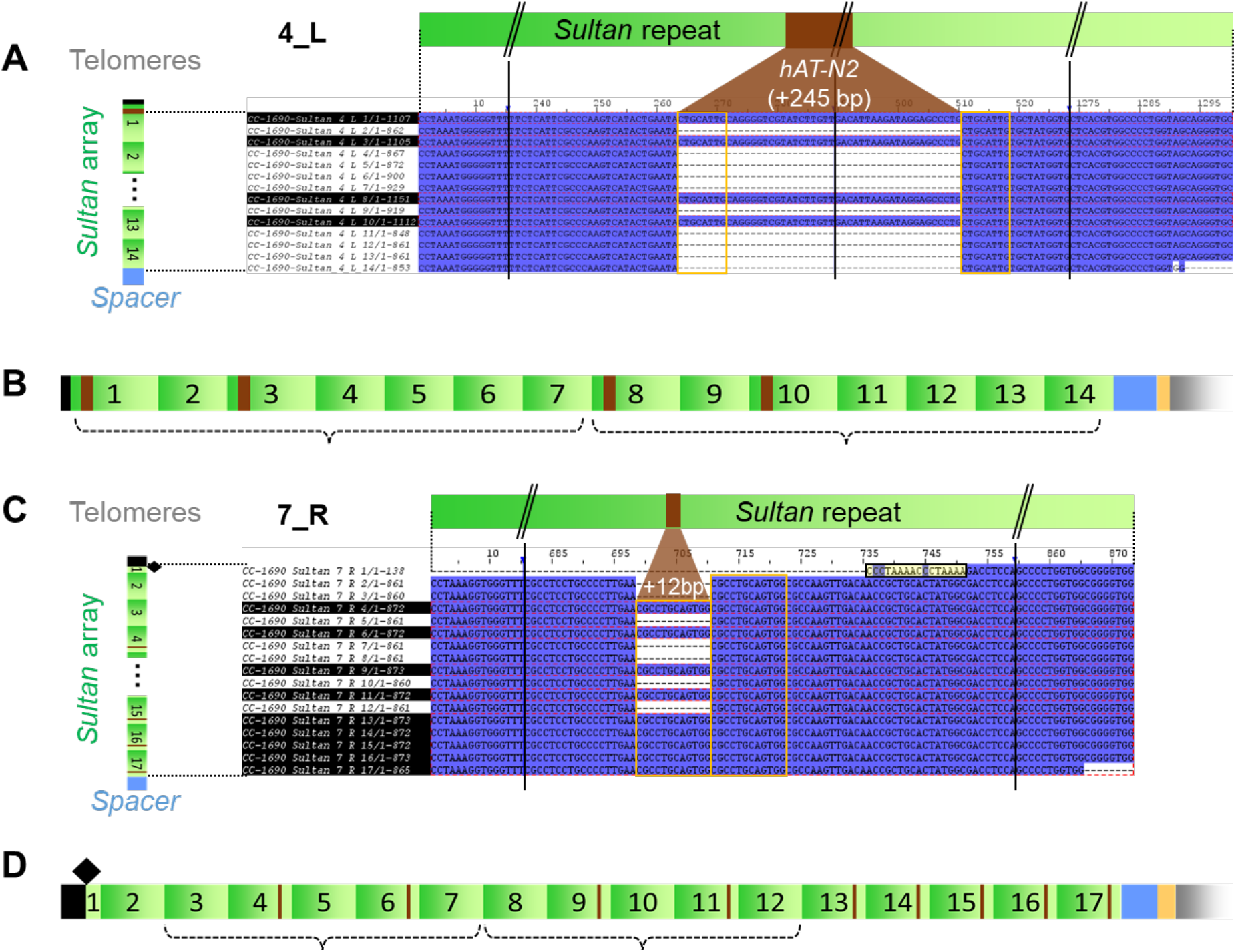
Evidence for multiple-copy duplication events in *Sultan* arrays. *(A)* and *(C)* Alignment of *Sultan* repeat sequences from subtelomeres 4_L and 7_R, from the telomere-proximal (top) to the *Spacer*-proximal (bottom). Conservation of nucleotide across repeats is indicated in dark blue. Vertical black bars and “//” denote sequence portions not shown. *Sultan* repeats highlighted in black present a large insertion, represented in brown. Orange frames highlight duplications (including a putative *hAT-N2* 8-bp target-site duplication in 4_L). In *(C)*, the end of the telomere is highlighted in yellow. *(B)* and *(D)* Sketch of 4_L and 7_R *Sultan* arrays with the conserved insertions (brown). Dotted brackets indicate multi-*Sultan* segments likely duplicated “en bloc” on each subtelomere.

### Four subtelomeres display distinct repeat arrays composed of *Subtile* and *Suber* elements

As depicted in Fig. 1, *Sultan* arrays were not adjacent to telomeres in class C subtelomeres 3_R, 5_R, 15_L and 16_L. Using repeat detectors XSTREAM and TRF, we found the sequence of variable length on the telomeric side to contain two new types of repeats described below, unrelated to the *Sultan* element, as well as vast low-complexity regions and short repeats.

All class C subtelomeres contained a ~190 bp repeat that we named *Subtile* for *SUBTelomeric repeat of Intermediate LEngth* (Fig. 4A, B and D). The 133 *Subtile* copies found in the CC-1690 assembly formed 29 tandem repeat arrays of various lengths, each containing between 1 and 12 *Subtile* copies. Several types of INDELs were detected upon alignment of *Subtile* copies (Fig. 4A; Supplemental File F1). For example, the last copy of an array on the centromere side was always truncated at the 3’ end side, by either 47 nt (Fig. 4A, blue) or 57 nt (Fig. 4A, dark green); in 6 arrays, the telomere-proximal copy was 5’-truncated by 134 nt (Fig. 4A, red); in 6 other arrays, the 6^th^ copy was larger due to an unrelated extra sequence of 146 nt (Fig. 4A, orange). Various combinations of these variants create 6 main types of *Subtile* arrays (Fig. 4B) and the dotted lines suggest possible routes for their generation. The number of arrays per subtelomere also varied greatly, from 1 (5_R, 16_L) to 21 (3_R). The structural alignment of the 29 arrays (Fig. 4D, simplified as plain boxes as shown in Fig. 4B) and their pattern of localisation in the subtelomeres suggested that full arrays and even series of arrays duplicated and propagated between chromosome arms. The analysis of non-repetitive sequences found between the arrays (“Other DNA” in Fig. 4D) suggested that they were likely duplicated along with the *Subtile* arrays.

**Figure 4.**
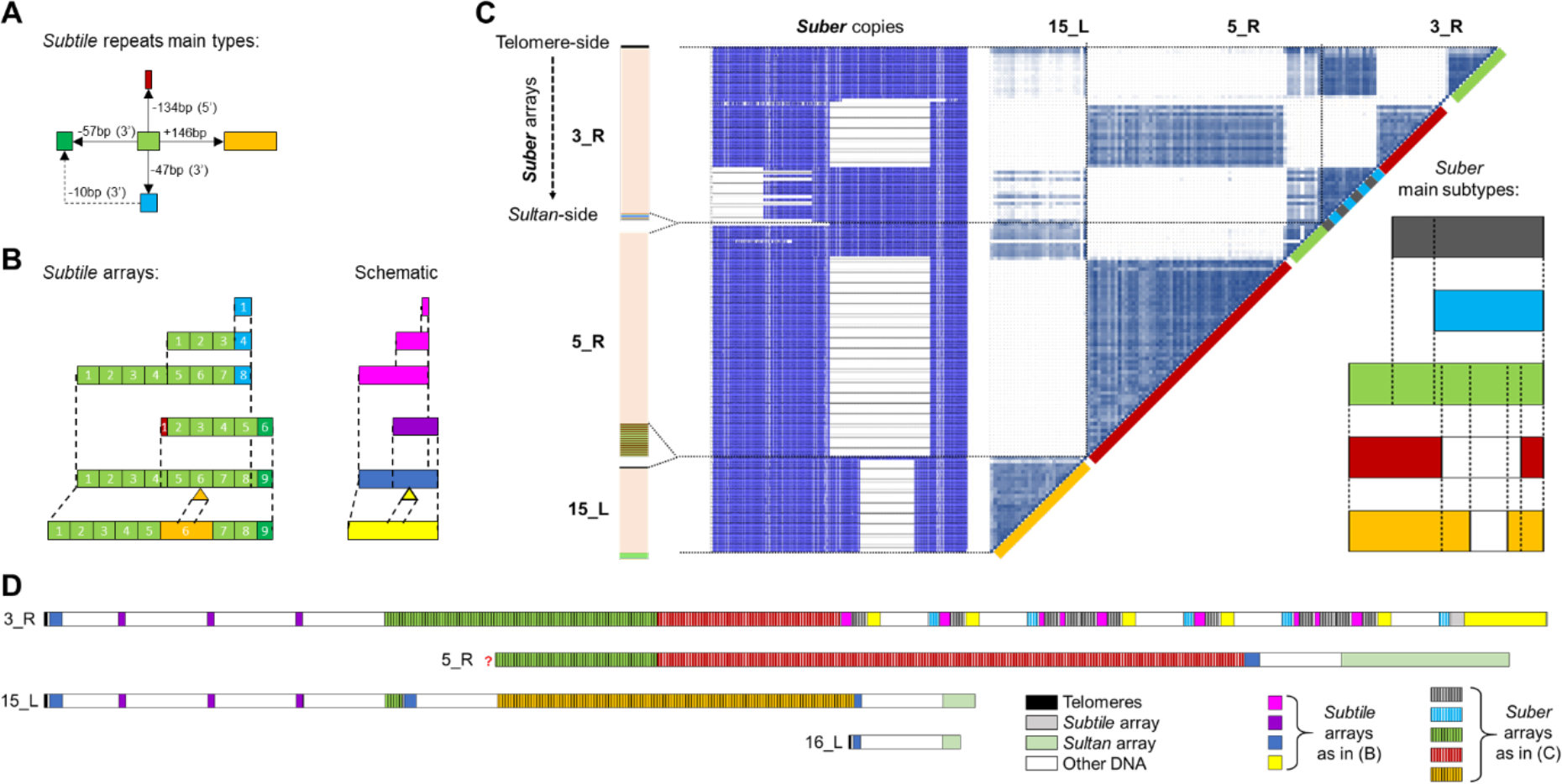
Arrays of *Subtile* and *Suber* repeats populate four subtelomeres in CC-1690. *(A)* Diagram of putative insertion/deletion steps depicting *Subtile* repeat variants, based on sequence alignment (Supplemental File F1). *(B)* Diagram of structural variations in *Subtile* arrays (left) and their simplified plain box representation (right). *(C)* Alignment (Supplemental File F1) and distance matrix of all *Suber* repeats. Light to dark blue color scale indicates increasing conservation. Subtypes colored on the diagonal of the distance matrix are depicted in the lower right diagram. *(D)* Map of *Subtile* and *Suber* arrays in class C subtelomeres drawn at scale following color code shown in *(B) and (C)*. Telomeres are shown as black boxes, *Sultan* repeat arrays as pale green boxes, other DNA sequences in white.

We further identified a third type of repeat in the 5_R, 3_R and 15_L subtelomeres, up to 2450 bp in length, that we called *Suber* for *SUBtelomeric Extra-long Repeats*. The 147 *Subers* assembled into massive arrays and analysis of the *Suber* variants indicated that they were also generated by segmental duplication. Four large INDELs (> 400 bp) allowed us to define five main types of *Suber* (Fig. 4C; Supplemental File F1), which formed a homogeneous array on subtelomere 15_L (52 kb) and two similar hybrid arrays on subtelomeres 5_R and 3_R (108 kb and 66 kb respectively) (Fig. 4D). In addition, subtelomere 3_R carried individual *Suber* copies between the *Subtile* arrays found in the centromere-proximal region. As for *Subtile* repeats, the similarity between subtelomeres 5_R and 3_R indicated an inter-chromosomal recombination, but the different numbers of 2461-bp *Suber* copies (green, 10 in 5_R vs. 16 in 3_R) and 1475-bp *Suber* copies (red, 68 vs. 35), suggested that *Suber* repeats continued to propagate *in situ* after the recombination event. In publicly available RNA sequencing datasets, we found evidence that *Suber* repeats might be transcribed. Moreover, BLAST and Conserved Domain searches indicate homology with bacterial HNH endonucleases, which belong to the homing endonuclease superfamily and can code for self-splicing introns and inteins (http://pfam.xfam.org/family/HNH). Only a few *Subers* contained these HNH-like regions in putative open-reading frames. *Suber* arrays also contained telomere-like sequences, corresponding to up to 10 degenerated repeats at the junction between *Suber* elements.

### Transposable elements populate subtelomeres downstream of the *Spacer* sequence

TEs are quite common in the subtelomeres of many organisms and can even function as telomeres in *D. melanogaster* (Pardue and DeBaryshe 2011). In *C. reinhardtii*, we found that, downstream of the G-rich repeats, a region most often spanning 5 to 15kb but reaching ~50 kb on some chromosome arms was generally populated by TEs, with exon density increasing progressively beyond these regions towards the centromeres (Supplemental Fig. S6). More specifically, the *L1* LINE element *L1-5_cRei* was found to specifically target the (GGGA)_n_ motif (Supplemental Fig. S4) and its copy number was enriched more than 50 fold in the 20 kb immediately downstream of *Spacers* relative to the rest of the genome. It is possible that *L1-5_cRei* has evolved such a targeted insertion sequence as a result of the abundance of the G-rich repeat in subtelomeres, which may serve as a safe haven that minimizes any deleterious effects of insertion.

### Interstrain variations provide insights into subtelomere evolution

To investigate the evolution of subtelomeres in *C. reinhardtii*, we compared the *Sultan* repeats in the CC-1690 genome with those in two other commonly used *C. reinhardtii* strains and one wild isolate, for which assemblies were generated from PacBio sequencing data and will be released in the near future (Craig et al., *in prep*). Compared with the genome assembled from Nanopore data, those obtained by PacBio were more often truncated at chromosome extremities, probably due to shorter read lengths, resulting in a smaller number of ends with telomeric repeats. Nevertheless, most *Sultan* arrays were at least partially assembled.

Among the strains studied, CC-503 has served as the long-term strain for the reference genome (Merchant et al. 2007), while CC-4532 (which was derived from a cross of CC-1690 and CC-124) has been assembled as a mating type *minus* strain as part of the forthcoming reference genome update. In contrast to these laboratory strains, CC-2931 is a wild isolate from North Carolina (USA), highly genetically differentiated from the other three (Flowers et al. 2015; Craig et al. 2019). As a result of the presence of two divergent genomes in their ancestry and of various subsequent crosses, all laboratory strain genomes consist of a mosaic of two alternative haplotypes (Gallaher et al. 2015). The minor haplotype, known as haplotype 2, covers a maximum of 25% of the genome and is expected to affect 8 chromosome ends (6_L, 8_L, 9_L, 10_R, 12_L, 12_R, 16_L, 16_R)(Gallaher et al. 2015). As the two haplotypes were inherited from a single isolated zygospore, genetic differences between them are expected to reflect diversity at the population level.

*Sultan* repeat consensus sequences from each subtelomere in all four strains were aligned to generate a phylogenetic tree using the maximum likelihood substitution algorithm (Fig. 5). When the sequence data were available, the *Sultan* repeats of the same chromosome end but from the different laboratory strains often grouped together, suggesting that most *Sultan* arrays have not relocated since these strains were genetically separated in the laboratory. A notable exception is the grouping of CC-503 2_R with 9_L in other strains, due to a documented reciprocal translocation between these chromosome arms that occurred in the laboratory history of CC-503 (Craig et al., *in prep*). In addition, CC-4532 carries haplotype 2 at 6_L (Gallaher et al. 2015), alternative to the haplotype of CC-1690 and CC-503, and we found that its *Sultan* consensus sequence at this subtelomere was highly similar to that found on 11_L in all three laboratory strains. *Sultan* repeats were much more divergent in the wild isolate CC-2931: a clear grouping with laboratory strains was observed only for 9 out of 24 available extremities, suggesting that allelic variation in subtelomeres is more pronounced at the species-wide level than the population level. In particular, CC-2931 6_L grouped with 10_R in other strains, not 11_L as in haplotype 2, thus representing a different allele.

**Figure 5.**
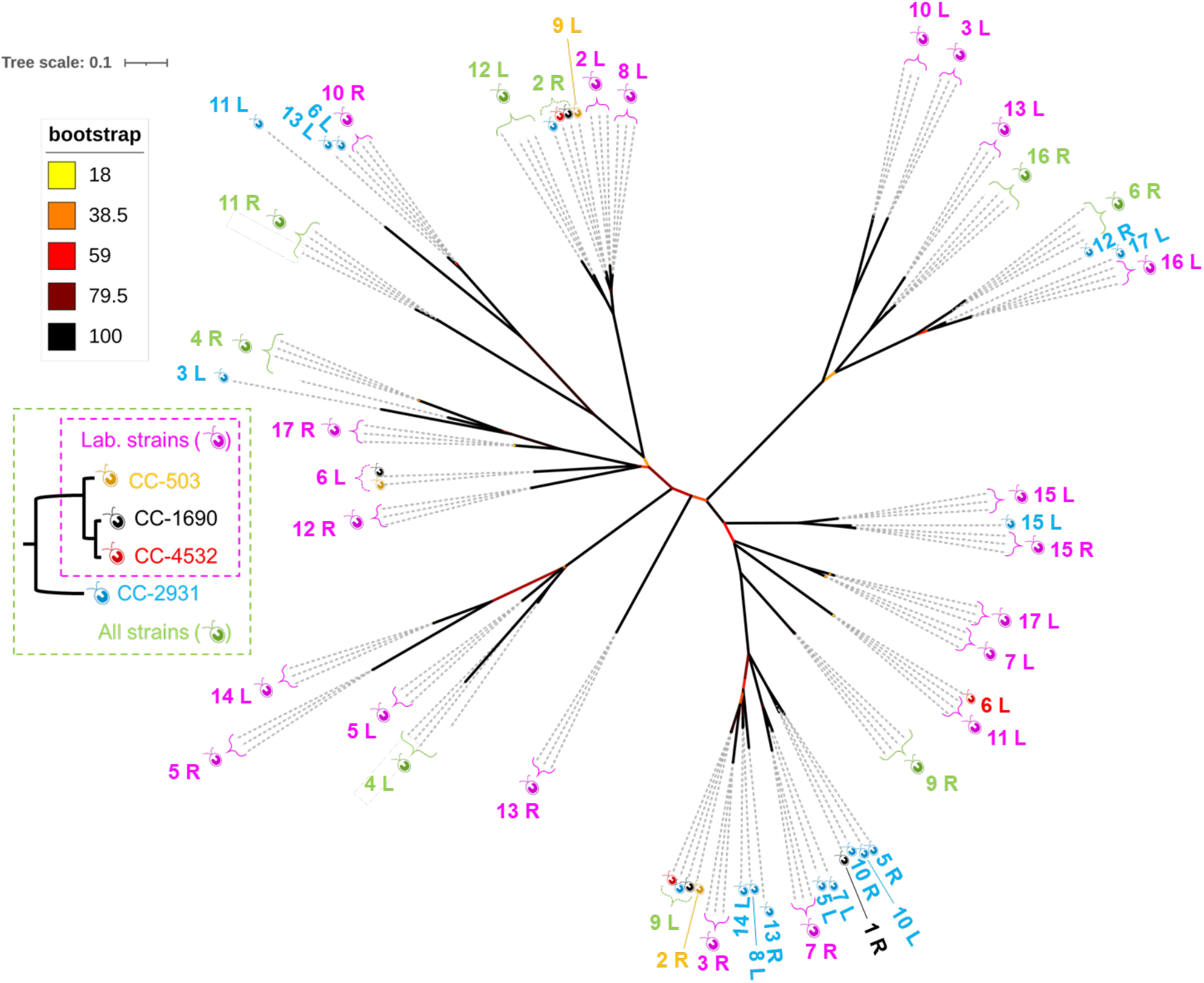
Phylogenetic tree of *Sultan* repeats in three laboratory strains and a wild isolate. *Sultan* consensus sequences from each chromosome end (Supplemental File F1) were aligned to generate a maximum-likelihood unrooted phylogenetic tree. Branch length and colour respectively represent substitution rates relative to the tree scale and bootstrap value (from the lowest in yellow to 100% in black). Chromosome ends clustering as closest homologs in all strains or in laboratory strains are grouped as green or pink symbols, respectively, with individual strains displayed in color for more complex groupings.

Interestingly, we found that the phylogenetic tree of the *Spacer* sequences from different subtelomeres was poorly concordant with the phylogeny of the *Sultan* element, as shown for CC-1690 (Supplemental Fig. S7), which might indicate that the *Spacers* mutated at a faster rate.

To investigate the variability in the number of copies in a given *Sultan* array between strains, we mapped Illumina sequencing reads of laboratory strains (Flowers et al. 2015) against the subtelomere-specific *Sultan* consensus sequences from CC-1690. We normalized the median nucleotide coverage of each *Sultan* consensus by the average whole genome coverage (Fig. 6). As a control, plotting the results from CC-1690 Illumina sequencing against the number of *Sultan* repeats observed in the CC-1690 end-to-end chromosomal assembly (Fig. 6A, blue) showed a linear relationship with only a slight overestimation of the repeat number. The same approach was then applied to CC-503 (Fig. 6A, orange) and other strains. The overall distribution of *Sultan* copy number across strains is shown as a boxplot of repeat counts for each subtelomere consensus (Fig. 6B) and a detailed comparison is displayed in Supplemental Fig. S8. Repeat counts were generally close to that of CC-1690 for most subtelomeres (median CV = 20%). Several of the major differences were in agreement with the expected distribution of the two alternative haplotypes amongst strains (Gallaher et al. 2015): CC-1009 and CC-408 had shorter subtelomeres at 6_L, 9_L and 12_R, in accordance with their carrying haplotype 2 at these loci. The shorter 6_L and 12_R subtelomeres were also found in CC-124 (also haplotype 2); at 8_L, strains with haplotype 1 (CC-503, CC-125, CC-1009, CC-408) had longer arrays than those with haplotype 2 (CC-1690, CC-1010), except for CC-124. For *Suber* and *Subtile* repeats, we found mapping Illumina reads datasets of all laboratory strains, but only in some of the available wild isolates, indicating that they might not be fully conserved in the species.

**Figure 6.**
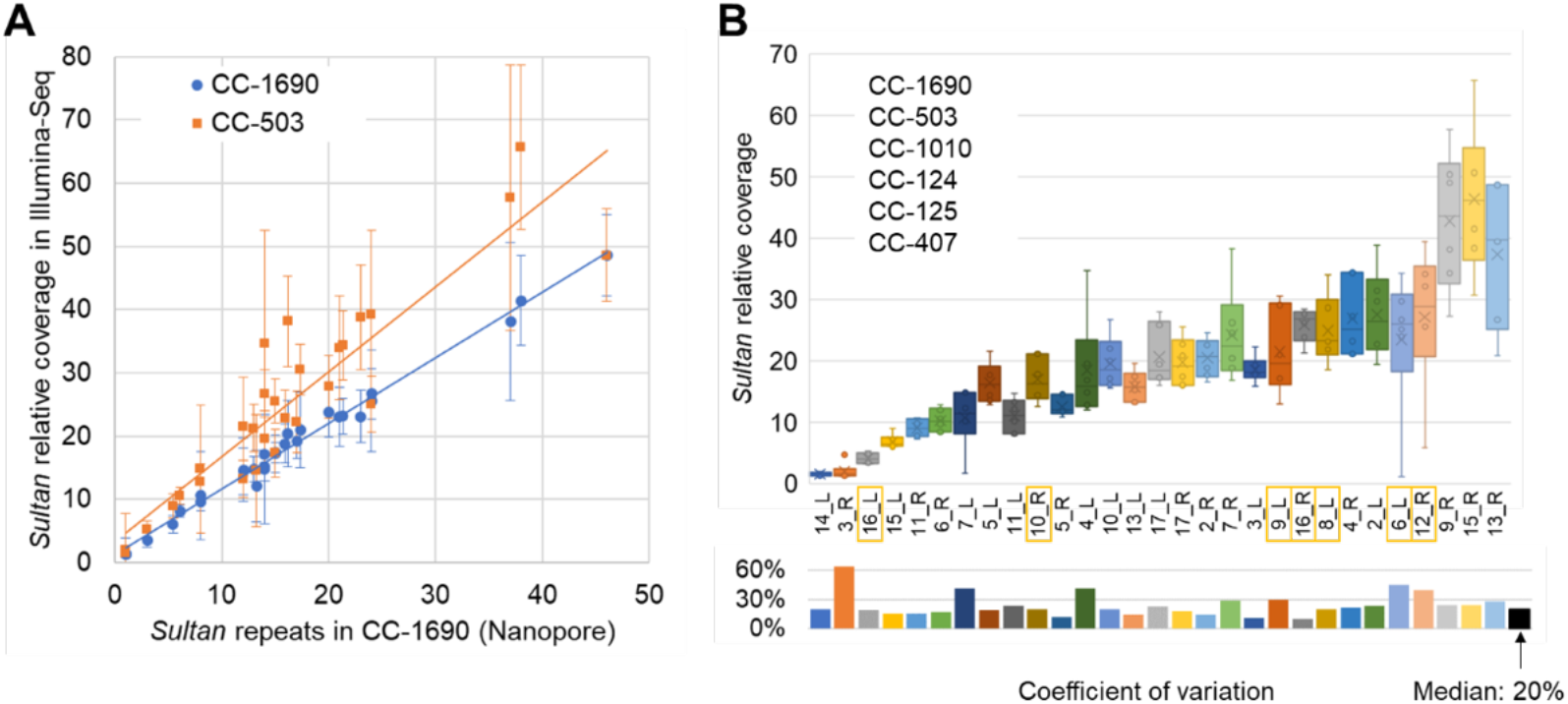
Count of *Sultan* repeats in distinct laboratory strains. Deep sequencing data were mapped to genome assembly of laboratory strain CC-1690 (Supplemental File F3). Estimates of *Sultan* repeat count in each subtelomere are calculated from the median read depth of *Sultan* consensus sequences. *(A)* Plot of *Sultan* repeat count estimates for CC-1690 (blue) and CC-503 (orange) against the actual repeat count observed in Nanopore sequencing of CC-1690. Shown are the median depth (±SD) and trend lines using the least-squares method. *(B)* Boxplot distribution (top) and coefficients of variation (bottom) of repeat count in laboratory strains for each subtelomere (see Supplemental Fig. S8 for strain-specific count, including more distant laboratory strains). Subtelomeres potentially affected by the distribution of haplotype blocks among these strains are highlighted.

### Subtelomeres in other green algae

We wondered whether a subtelomeric organization similar to that in *C. reinhardtii* would be found in other algae. We concentrated on the few algal genomes that present the degree of completeness and accuracy that was needed for this analysis. The closest known relatives of *C. reinhardtii* are *C. incerta* and *C. schloesseri*, for which highly contiguous long read genome assemblies were recently produced (Craig et al. 2020). They show a high degree of synteny with *C. reinhardtii* (84% and 83% of their genome length, respectively). Several chromosomes (6 and 4, and possibly others) appear almost fully conserved with *C. reinhardtii*, and they putatively share a centromeric structure based on arrays of *Zepp*-like retrotransposons. We were therefore surprised to find by Blast no trace of any of the *Sultan*, *Subtile* or *Suber* repeats described in *C. reinhardtii*. In *C. incerta*, 4 of the 5 contigs showing terminal arrays of the 8-bp telomeric repeats shared a well-conserved 350 nt repeat forming immediately-subtelomeric arrays (Fig. 7). We called this repeat *Subrin*, for *SUB*telomeric *R*epeat of *C. INcerta*. *Subrin* arrays were found in 29 additional contigs lacking telomeres, but in an orientation generally consistent with a subtelomeric position. Some arrays were very extensive and we counted a total of 1819 *Subrin* copies in the assembly. *Subrin* copies were more similar within an array than across arrays, again indicating preferential local tandem duplications. In 29 cases, we could collect the sequence immediately upstream of the *Subrin* array, and found that 24 of them started with a homologous spacer sequence, generally spanning ~1.2 kb. No G-rich repeat region was observed. We conclude that in *C. incerta* also, the majority of the chromosomes comprise a repetitive subtelomeric sequence anchored on a conserved spacer, even though the sequences themselves were unrelated to those in *C. reinhardtii*.

**Figure 7.**
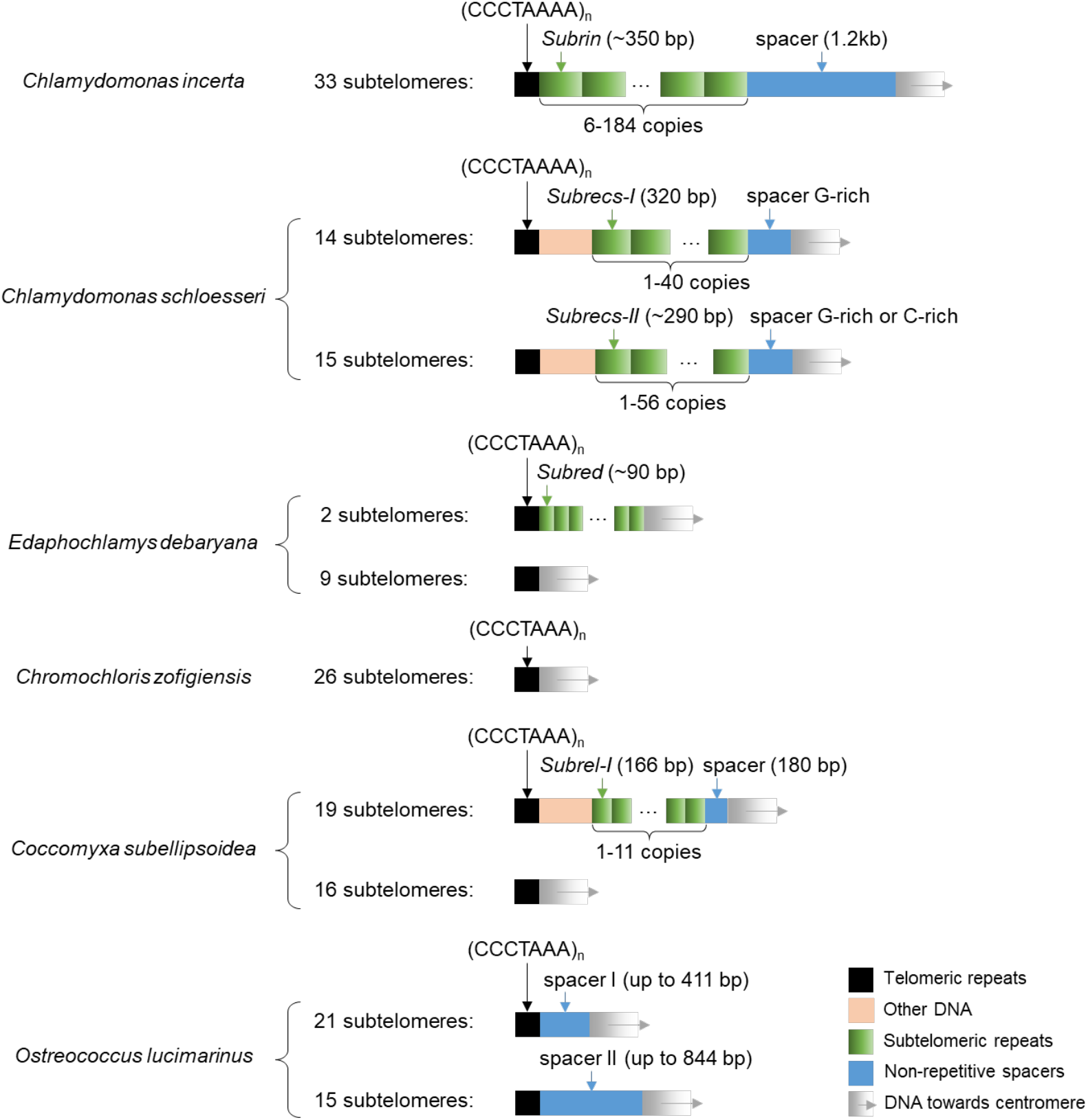
Subtelomere architectures in microalgae. Genome assemblies of the indicated species were searched for repeats (green boxes) and shared (blue) sequences near telomeres (black box). Intervening DNA and downstream chromosome arms are shown in pink and grey, respectively.

Subjected to the same analysis, *C. schloesseri* revealed a similar type of subtelomere organization (Fig. 7), but again based on repeats unrelated to either the *Sultan* or the *Subrin*. We called these repeats *Subrecs*, for *SUB*telomeric *RE*peat of *C. Schloesseri*. However, they displayed more heterogeneity than in *C. reinhardtii* or *C. incerta*. We distinguished two types, unrelated in sequence, called *Subrecs-I* (319-321 bp, 233 copies) and *Subrecs-II* (266-327 bp, 298 copies). They formed arrays in respectively 14 and 15 contigs but immediately adjoined the terminal telomeric repeats only in respectively 2 and 6 cases. This is because many contigs carried one or even two non-terminal telomeric repeat array, sometimes in addition to a terminal one. Internal telomeric arrays were often adjoined by or embedded in a *Subrecs* array. Noticeably, a contig carried only type-I or type-II *Subrecs*, never a mixture, suggesting a history of subtelomeres in *C. schloesseri* with complex recombination processes involving mostly *cis*-sequences. As in *C. reinhardtii*, the centromere-proximal *Subrecs* were adjacent to a conserved spacer, again with a short 3’-truncation (2 nt for *Subrecs-I*, 7 for *Subrecs-II*). In terms of sequence homology, the spacers themselves were of two types: type G (G-rich) was associated with *Subrecs-I* or -*II* arrays, type C exclusively with *Subrecs-II*.

*Edaphochlamys debaryana* is a more distant relative of *C. reinhardtii*, but also groups within the core-*Reinhardtinia* clade of the Chlamydomonadacae. The synteny with *C. reinhardtii* is less marked (46%) and the assembly is less contiguous. Here telomeric repeats of 7 nt (CCCTAAA) were observed, but in only two telomeres were they associated with a subtelomere-specific repeat which we called *Subred* (*SUB*telomeric *R*epeat of *Edaphochlamys Debaryana*) (Fig. 7). At a further phylogenetic distance, the genome of *Chromochloris zofingiensis*, a Chlorophycea of the class Sphaeropleales, also showed 7-nt telomeric repeats but an absence of subtelomere-specific repeats.

Repetitive subtelomeres can also be found in other green algae. In the almost fully assembled genome of the Trebouxiophyceae *Coccomyxa subellipsoidea*, the 20 chromosomes carry 7-nt telomere repeats at both extremities. In 19 extremities, the subtelomere comprises what we called a *Subrel-I* repeat (*SUB*telomeric *R*epeat of *Coccomyxa subELlipsoidea*) of 166 nt (1 to 11 copies per extremity, 86 in total) (Fig. 7). Only in 3 cases was the array adjacent to the telomere. Again, a conserved spacer sequence of ~180 nt was found on the centromeric side of every *Subrel* array, and in one case 5’-truncated and abutting the telomeric array, suggestive of a deletion of the *Subrel* array. In addition, other repeats called *Subrel-II* (~90 nt) and *Subrel-III* (~19 nt) were found in respectively 3 and 2 subtelomeres.

In the Mamiellophyceae *Ostreococcus lucimarinus*, with 21 chromosomes, no subtelomeric repeat could be identified. However, many extremities shared a homologous sequence immediately after the telomere (Fig. 7). Type-I (up to 411 nt) was found in 21 extremities, Type-II (up to 844 nt) in 15. In both groups, especially Type-II, some subtelomeres were truncated at the 5’ end and the junction with the telomeric repeat was in various phases. Combined with the presence of fragments of the Type-I sequence at the 5’ of a Type-II, this suggests a history of partial deletions and repair.

## Discussion

### A comprehensive description of the architecture of subtelomeres in *C. reinhardtii*

Subtelomeres are notoriously difficult to assemble due to their repetitive nature. Previous reference genomes of *C. reinhardtii* failed to provide a clear picture of the subtelomeres and also lacked telomere sequences at most extremities. Using long read sequencing data (PacBio and Oxford Nanopore Technology) and *de novo* genome assemblies (Liu et al. 2019; Craig et al. 2020; O'Donnell et al. 2020) (Craig et al., *in prep*), we now provide a nearly complete map of all chromosome extremities in *C. reinhardtii*, including telomere sequences at 31 out of 34 extremities. Given the mean read length and *N50*, both equal to 55 kb, of the Nanopore reads, and the contiguity of our assembly, we are confident that our description of the subtelomeres is accurate, especially for the exact number of repeated elements in each subtelomere.

We describe three new types of repeated elements present in *C. reinhardtii* subtelomeres. The *Sultan* element is the most abundant, found in 31 out of 34 subtelomeres, absent from the rest of the genome and therefore can be considered as specific of a canonical subtelomere. We classify the subtelomeres into four groups based on their organization. The most common, class A, corresponds to the following architecture: telomere sequences, array of *Sultan* repeats, *Spacer* sequence, G-rich microsatellite and TEs. The length between the telomere and the microsatellite in this class is typically 10-30 kb. Class B subtelomeres are similar except that their *Sultan* elements contain large insertions of 250-1000 bp. The four class C subtelomeres contain other repeated sequences, called *Subtile* and *Suber*, between the telomeres and the *Sultan* elements, and can be much longer (*e.g.* > 200 kb for 3_R). Finally, three chromosome extremities contain ribosomal DNA (1_L, 8_R and 14_R) and none of the repeated elements found in the other classes: they were grouped in class D. In one of them (1_L) we identified telomere sequences adjacent to the rDNA, capping the extremity, showing that rDNA sequences can constitute a subtelomere by themselves. The repetitive nature and the expected size of the rDNA sequences in subtelomeres 8_R and 14_R made it impossible for the assembly to reach telomere sequences at these two extremities. The subtelomeric localization of rDNA repeats in *C. reinhardtii* is reminiscent of rDNA sequences found at some chromosome extremities in *A. thaliana* (Arabidopsis Genome 2000), in some species of the *Allium* genus (Pich and Schubert 1998; Fajkus et al. 2016), and in *Schizosaccharomyces pombe* (Wood et al. 2002). It is possible that the heterochromatic nature of telomeres/subtelomeres and rDNA makes their proximity an advantageous feature for the genome, as suggested by heterochromatin assemblies acting functionally as telomeres in some telomerase-negative *S. pombe* survivors (Jain et al. 2010). Whether *Sultan*, *Subtile* and *Suber* repeat arrays can form heterochromatin remains to be investigated, but this possibility might explain their presence at subtelomeres.

The three repeated elements we describe (*Sultan*, *Subtile* and *Suber*) are uniquely found at subtelomeres. We do however find some homology between the central part of the *Sultan* sequence (nt ~170-510) and the centromere-associated minisatellite *MSAT2_CR*, between sequences inserted in some *Sultan* elements and TEs, and between the *Suber* element and the HNH endonuclease domain superfamily. Besides, the *Suber* elements and the *Spacer* sequences are transcribed. These putatively non-coding but spliced and polyadenylated transcripts are similar to sub-TERRA and other subtelomeric transcripts, as described in multiple organisms (Azzalin and Lingner 2015; Kwapisz and Morillon 2020), with potential functions in telomere maintenance that remain to be investigated. The 5’ part of the *Spacer* element functions as a promoter, active essentially at dusk and during the first phase of night in a light-dark cycle, concomitantly with replication and histone deposition (Strenkert et al. 2019).

### Molecular mechanisms of segmental duplication and contraction

An important finding is that *Sultan* elements show higher similarity within a subtelomere than between subtelomeres, suggesting a very low frequency of rearrangements involving different extremities. This observation is consistent with the relatively low efficiency of homology-based recombination in vegetative *C. reinhardtii* cells (Zorin et al. 2005). It is however in contrast with what is known in other species where subtelomeric regions show signatures of frequent interchromosomal recombination between repeated sequences (Louis et al. 1994; Linardopoulou et al. 2005; Chen et al. 2018). Nevertheless, although infrequent, rearrangements between subtelomeres did occur as evidenced by the propagation of the *Sultan* elements on most subtelomeres and the similarities between the arrangement of *Subtile* and *Suber* repeats in different subtelomeres. The high similarity between *Sultan* elements belonging to the same subtelomere suggests that at some point, only one *Sultan* element was present at a given subtelomere, or maybe sometimes two for the *Sultan* arrays composed of two slightly different types of *Sultan* (e.g., 4_L or 7_R, Fig. 3). Another argument for this possibility is that *Sultan* elements in each class B subtelomere contained a single type of insertion. Alternatively, frequent intra-subtelomere gene conversion or other recombination-based mechanism events might homogenize the sequence of the *Sultan* elements within a subtelomere. We therefore propose that either (i) a single ancestral *Sultan* element (or possibly two) colonized each subtelomere, diverged from each other and underwent multiple segmental duplications *in cis*, or (ii) *Sultan* arrays colonized different subtelomeres, diverged and collapsed to only one copy per subtelomere (or possibly two), which then duplicated *in cis*, or (iii) *Sultan* arrays colonized different subtelomeres and underwent homogenization within each subtelomere.

Contraction events might be promoted by the seed telomere sequence present at the 5’ end of the *Sultan* element. Indeed, since telomeres and repeated elements are difficult to replicate, DNA breaks at a subtelomere due to a replication defect might be repaired by telomere healing primed by the seed sequence. This possibility is supported by the fact that the telomere seed sequence of the *Sultan* closest to the telomere is in phase with and transitions seamlessly into the telomeric tract in most cases. Such a mechanism would lead to the terminal deletion of a variable number of *Sultan* elements. In 9 subtelomeres, however, the transition to the telomeric repeat occurs within a *Sultan* copy (diamonds in Fig. 1), at various phases of the telomeric repeat and in only one case at an internal telomere-like sequence of the *Sultan*. Here, a double strand break and NHEJ using a telomere fragment could account for the observed repaired structure.

Several mechanisms can explain the segmental duplication of one or several tandem *Sultan* elements along a subtelomere: unequal SCE, rolling circles, replication slippage, BIR, or HR. Because of the greater similarity within *Sultan* elements from the same subtelomere compared to other subtelomeres, we favor mechanisms that do not require other chromosome ends. We thus speculate that both expansion and collapse of *Sultan* elements have contributed to the current architecture of subtelomeres.

### Evolution of subtelomeres within *C. reinhardtii* and beyond

To provide an idea of how dynamic the subtelomeres of *C. reinhardtii* are, we compared the sequences of the *Sultan* and *Spacer* elements, as well as the copy number of the *Sultan* elements in each subtelomere, in different strains, including laboratory strains and a wild isolate. Based on PacBio-sequencing-based assemblies for two additional laboratory strains, we found a good conservation of the sequences of the *Sultan* elements with no evidence for subtelomere-specific rearrangements within the laboratory strains. Since Illumina sequencing data were available for a number of strains, we developed a method using the number of reads mapping to a consensus sequence for the *Sultan* elements in a given subtelomere to estimate their copy number in each subtelomere, without the need for a genome assembly. This showed that the copy number was in general well conserved with a few exceptions, which in almost all cases could be traced to strains carrying a distinct ancestral haplotype (Gallaher et al. 2015). We also analyzed a slightly less complete assembly of the wild isolate CC-2931, and observed a mixed pattern, with both conserved subtelomeres and evidence for polymorphic alleles, which may be created by the translocation of *Sultan* elements between chromosomes. Overall, subtelomere architecture appears to have undergone little evolution since introduction in the laboratory, but substantial polymorphism exists at the population and species-wide level, possibly paving a way towards speciation.

At a larger evolutionary scale, we found specific repeated elements for most species of green algae we looked into. Interestingly, subtelomere organization in these algae seemed to follow a structure similar to *C. reinhardtii*, with an array of repeated elements adjoining the telomere and a spacer sequence conserved across subtelomeres, but the repeated element and the spacer sequence were unrelated across species. This study was limited by the small number of chromosome level assemblies for green algae, but it suggests that subtelomeres in many green algae have converged to strikingly similar structures, with conserved species-specific (usually repeated) elements populating the subtelomere. Investigating the underlying properties that drove the propagation of these elements, such as hetechromatin formation, binding of particular factors, or transcription from the spacer sequences, might contribute to better understand subtelomere functions and evolution.

## Material & methods

### Genome assemblies and repeats

The genome assemblies for *C. incerta*, *C. schloesseri* and *E. debaryana* are described in (Craig et al. 2020). The CC-4532 and CC-503 (v6) assemblies are forthcoming and will be made available in the near future (Craig et al., *in prep*), that from CC-2931 was obtained by assembly of PacBio reads. For strain CC-1690 (“21gr”), recently released Nanopore raw sequencing data (Liu et al. 2019) were base-called and *de novo* assembled into chromosomes as described in (O'Donnell et al. 2020)(GenBank accession: JABWPN000000000). For the present work, we used a version prior to the Illimuna polishing step and used linkage data (Ozawa et al. 2020) to further scaffold the last unplaced contig (unplaced_1) to the end of chromosome 15, forming its right arm. Compared to our released genome (O'Donnell et al. 2020), we corrected a mistake in the assembly of subtelomere 9_R (a replacement contig is appended to the genome), which was distorted at the telomere-proximal side of the *Sultan* array by reads from 15_R. To do this, reads were first mapped against the whole genome using minimap2 (Li 2018), then extracted if they mapped to the 9_R and contained a mapping quality of 60. This subsample of reads was then used for re-assembly with Canu (V2) using default settings. Additionally, the 1_R end, which did not contain a telomeric sequence nor *Sultan* repeats at its apparent terminus, was analyzed by read mapping and we were able to recover a few reads, extending beyond the assembly and containing both telomere sequences and 14 *Sultan* repeats.

We extracted and analyzed the first 30 kb of the chromosome ends (300 kb for class C subtelomeres). Sequences from the right extremities were reverse complemented, so that both left and right chromosome ends started with telomeric repeats in the form of 5’-(CCCTAAAA)_n_-3’ tracts. Our numbering reads from telomere towards centromere.

Chromosome ends from CC-503, CC-4532 and CC-2931 were extracted from PacBio-sequencing-based assemblies. A notable difference in CC-503 is the translocation between chromosome arms 2_R and 9_L as compared to other laboratory strains. Genome coordinates and sequences were extracted using blast or seqret to generate gff and fasta files. A curated library of Volvocales TEs (Craig et al. 2020) was used to identify mobile and repetitive genetic elements, using RepeatMasker (http://repeatmasker.org/).

### Search for tandem repeats

We use the term “repeat” to refer to the finite pattern found in a repetitive sequence, “copy” to a specific instance of the repeat and “array” to a series of copies. Copies that are found in “tandem” in an array are in the same orientation not separated from each other by unrelated DNA sequences.

Sequences were analyzed using Tandem Repeat Finder (v4.04, parameters 3 5 5 80 20 100 2000) and X-STREAM (variable sequence tandem repeats extraction and architecture modeling, https://amnewmanlab.stanford.edu/xstream/) (Benson 1999; Newman and Cooper 2007). X-STREAM was run with default parameters, except “TR significance” was disabled and “Minimal word match” and “Minimum Consensus match” could be decreased down to 0.1 and increased up to 0.95 respectively, to allow detection of incomplete repeats at extremities of tandem arrays. Repeat consensus sequences were phased and used as blast queries to retrieve individual copies on the genome using EMBOSS (v6.4.0) seqret (Supplemental File F2). Multiple sequence alignments were generated with MAFFT (v7.130) with iterative refinement method G-INS-i. Pairwise distances were calculated using EMBOSS distmat with Jukes-Cantor substitution model.

Phylogenetic analyses and trees were generated using PhyML with generalized time-reversal (GTR) model for nucleotide evolution and drawn using Interactive Tree Of Life (https://itol.embl.de/). JAL-view (Waterhouse et al. 2009) and Bioedit (Hall 1999) were used for data visualization and calculation of consensus and logo sequences. Consensus sequences were computed from Advanced Consensus Maker (https://www.hiv.lanl.gov/cgi-bin/CONSENSUS_TOOL/consensus.cgi).

### Transcriptomics

Transcript dataset from (Strenkert et al. 2019) (accession number: GSE112394; strain CC-5390) was searched using each *Spacer* sequence as BLAST queries on NCBI server. Duplicate hits were discarded and coverage was calculated as total nucleotide amount.

Iso-Seq data (accession number: PRJNA670202; multiple laboratory strains) and ChIP-seq data (accession number: PRJNA681680; strain CC-5390) were used to assess transcription and H3K4me3 marks (Gallaher et al. 2021). Circular consensus sequence Iso-Seq reads were mapped against the CC-1690 assembly using minimap2 (parameters: -ax splice:hq --secondary no). ChIP-seq reads were mapped using bwa mem (Li 2013), duplicates were removed using the Picard tool MarkDuplicates (http://broadinstitute.github.io/picard/), and peaks were called with MACS v2 (parameters: callpeak - g 1.0e8 -B ----fix-bimodal --extsize 150) (Zhang et al. 2008).

### Genomic reads mapping

Illumina data for each strain (Supplemental Table ST3) were mapped against the whole genome of CC-1690 using bwa-mem (Li 2013). The bam file was used to calculate the average whole genome coverage and extract all reads mapping to *Sultan* arrays. This read subset was then aligned against all *Sultan* consensus sequences from the same strain. The fold increase in median coverage within each consensus, compared to the whole genome, was used as a measure of the number of repeats within each array from which the consensus was derived (Supplemental File F3).

## Acknowledgments

This work was supported by the Agence Nationale de la Recherche grants “AlgaTelo” (ANR-17-CE20-0002-01) and “PhenoVar” (ANR16-CE12-0019), by the “Initiative d'Excellence” program from the French State (Grant ‘DYNAMO’, ANR-11-LABX-0011-01), and by Ville de Paris (Programme Émergence(s)). RC is supported by a BBSRC EASTBIO Doctoral Training Partnership grant.

We thank the Department of Energy Joint Genome Institute and collaborators for pre-publication access to the genomes of *Chlamydomonas reinhardtii* strains CC-503 and CC-4532 in support of this analysis. We thank Maria Teresa Teixeira and Gilles Fischer for their critical reading of the manuscript.

