## Supplemental Information for "Architecture and evolution of subtelomeres in the unicellular green alga *Chlamydomonas reinhardtii*"

*: co-last authors

**Affiliations:**

[1]Sorbonne Université, CNRS, UMR7238, Institut de Biologie Paris-Seine, Laboratory of Computational and Quantitative Biology, 75005 Paris, France

[2]Institute of Evolutionary Biology, School of Biological Sciences, University of Edinburgh, EH9 3FL, Edinburgh, United Kingdom

[3]Sorbonne Université, CNRS, UMR7141, Institut de Biologie Physico-Chimique, Laboratory of Chloroplast Biology and Light-Sensing in Microalgae, 75005 Paris, France

| Chr. | Length | End | Telomere length | *Sultan* repeat | | Transition with... | | *Spacer* length | G-rich repeats | |
|---|---|---|---|---|---|---|---|---|---|---|
| | | | | Copy number | Length [b] | Telomeres [c] | *Spacer* [d] | | Consensus [e] | Length |
| 1 | 8131179 | _L | 113 | | None (rDNA) | | | | None | |
| | | _R[a] | 281 | 14 | 961 | Phased | Standard | 465 | GGGA | 780 |
| 2 | 8577918 | _L | 420 | 23 | 863 | Phased | Standard | 541 | GGGA | 394 |
| | | _R | 291 | 15.9 | 862 | 5'-526 | 432-3' | 564 | ggga | 137 |
| 3 | 9243487 | _L | 292 | 17 | 872 | Phased | Standard | 494 | GGGA | 453 |
| | | _R | 498 | 1 | 246 | 5'-246 | Standard | 525 | GGGA | 241 |
| 4 | 4015868 | _L | 125 | 14 | 862/1107 (245) | Phased | Standard | 536 | ggga | 165 |
| | | _R | 370 | 21.3 | 870 | 5'-596 | Standard | 1000 | GGGa | 82 |
| 5 | 3710367 | _L | 174 | 12 | 1157 (319) | Phased | Standard | 468 | GGGAGA | 426 |
| | | _R | n.d. | 13.2 | 1898 (1037) | Phased | 442-3' | 0 [f] | None | |
| 6 | 8901863 | _L | 313 | 24 | 868 | Phased | Standard | 470 | GGGA | 414 |
| | | _R | 70 | 8 | 879 | Phased | Standard | 524 | GGGA | 406 |
| 7 | 6402991 | _L | 350 | 8 | 844 | Phased | Standard | 476 | GGGAGA | 532 |
| | | _R | 348 | 16.2 | 873 | 5'-739 | Standard | 494 | GGGAGA | 247 |
| 8 | 4565225 | _L | 307 | 21 | 860 | Phased | Standard | 558 | ggga | 105 |
| | | _R | n.d. | | None (rDNA array) | | | | None | |
| 9 | 6779811 | _L | 50 | 17.3 | 725 | 5'-448 | Standard | 477 | GGGA | 216 |
| | | _R | 287 | 37 | 892 | Phased | Standard | 536 | GGGA | 472 |
| 10 | 6719514 | _L | 311 | 14 | 875 | Phased | Standard | 487 | GGGA | 523 |
| | | _R | 324 | 13 | 1138 (358) | Phased | Standard | 641 | GGGA | 369 |
| 11 | 4509270 | _L | 378 | 12 | 862 | Phased | Standard | 481 | GGcA | 539 |
| | | _R | 381 | 6.1 | 1402 (544) | 5'-1300 | Standard | 494 | GGcA | 137 |
| 12 | 9804983 | _L | 27 | 1.6 | 840 | Phased | 494-3' | 40 [f] | None | |
| | | _R | 122 | 24 | 844 | Phased | Standard | 499 | GGGA | 350 |
| 13 | 5293517 | _L | 305 | 14 | 1265 (387) | Phased | Standard | 529 | GGGA | 338 |
| | | _R | 394 | 46 | 873 | 5'-791 | 265-3' | 254 [g] | GGGA | 747 |
| 14 | 4233087 | _L | 380 | 1 | 1116 (263) | Phased | Standard | 496 | gGGGA | 196 |
| | | _R | n.d. | | None (rDNA array) | | | | None | |
| 15 | 3797459 | _L | 493 | 5.4 | 866 | 5'-491 | Standard | 497 | GGGA | 354 |
| | | _R | 313 | 38 | 879 | Phased | Standard | 494 | GGGA | 362 |
| 16 | 7887886 | _L | 371 | 3 | 871 | Phased | Standard | 538 | GGGA | 192 |
| | | _R | 347 | 20 | 872 | Phased | Standard | 541 | GGGA | 292 |
| 17 | 6886220 | _L | 136 | 15 | 848 | Phased | Standard | 476 | GGGA | 203 |
| | | _R | 188 | 15 | 862 | Phased | Standard | 504 | GGGA | 347 |
| | **Median:** | | **311** | **14.5** | **867** | | | **501** | | **349** |

**Supplemental Table ST1. Description of chromosome extremities in CC-1690.**

Chr: chromosome; All lengths expressed in bp; [a]: only from two reads; [b]: including extra DNA in brackets (not used to compute the median); [c] and [d]: positions of the first nucleotide of the 1st *Sultan* repeat and of the last nucleotide of the last *Sultan* repeat, respectively, with respect to the consensus; [e]: highly (uppercase) and moderately (lowercase) conserved nucleotides; [f]: sequence downstream not related to other *Spacer* sequences; [g]: *Spacer* from 13_R truncated on the *Sultan* repeat side; n.d.: not detected.

| Subtelomere (number of repeats) | Position within the *Sultan* elements | Type of insertion (Family) | Subject cover | Identity (%) |
|---|---|---|---|---|
| **4_L (1)** | 272-509 | *hAT-N2_cRei* (DNA/*hAT*) | 100% | 98% |
| **4_L (3)** | 272-507 | | | 97% |
| **4_L (8)** | 272-505 | | | 97% |
| **4_L (10)** | 272-509 | | | 96% |
| **5_L *** | 668-994 | *rnd-1_family-144* | 10% | 95% |
| **5_R *** | 666-1735 | | 100% | 97% |
| **9_R *** | 516-563 | *RTEX-1_cRei* (LINE/*RTEX*) | 1% | 100% |
| **10_R *** | 182-537 | *LTR-N1_cRei* (LTR) | 100% | 97% |
| **11_R *** | 334-389 | *Gypsy-20_cRei_LTR* (LTR/*Gypsy*) | 65% | 93% |
| | 403-874 | | | 93% |
| **13_L *** | 239-629 | *Chlamys-N1_cRei* (PLE/*Chlamys*) | 19% | 95% |
| **14_L** | 350-624 | *L1-7_cRei* (LINE/*L1*) | 9% | 88% |

**Supplemental Table ST2. Large insertions in *Sultan* repeats.**

*: Found in all *Sultan* repeats of the indicated subtelomere (only the statistics for the telomere-proximal repeat are then shown). The insertions for *RTEX-1_cRei*, *L1-7_cRei* and *Chlamys-N1_cRei* all consist of the 3′ end of the elements and are consistent with 5′ truncation. *hAT-N2_cRei* and *LTR-N1_cRei* are full-length. *Gypsy-20_cRei_LTR* is likely a solo LTR. *rnd-1_family-144* corresponds to a putative TE of unknown classification (*unknown-10_cRei*).

| Strain | SRA accession (Flowers et al. 2015) |
|---|---|
| CC-1690 | SRX 823854 |
| CC-503 | SRX823862 |
| CC-1010 | SRX823850 |
| CC-124 | SRX823851 |
| CC-125 | SRX823852 |
| CC-407 | SRX823859 |
| CC-1009 | SRX823849 |
| CC-408 | SRX823860 |

**Supplemental Table ST3. SRA accession numbers of Illumina-sequencing datasets (Flowers et al. 2015) used in this study for *Sultan* coverage.**

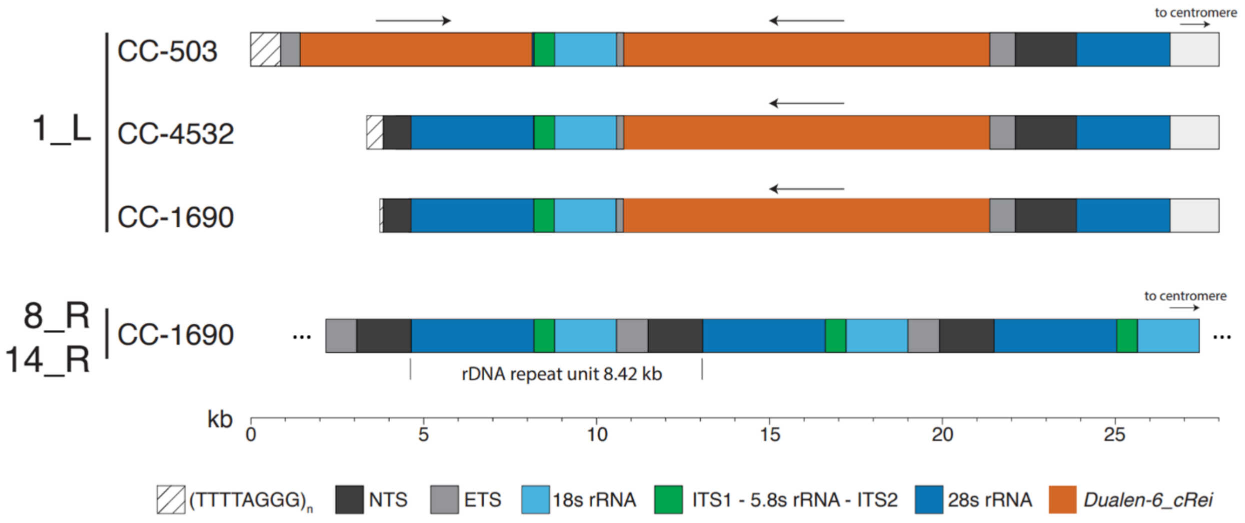


**Supplemental Figure S1. Schematic of the rDNA.**

Position of the distinct elements found at subtelomere 1_L as compared to 8_R and 14_R subtelomeres. Ribosomal DNA shown as blue, grey and green boxes; *Dualen-6_cRei* retrotransposon shown in orange with arrow indicating 3′ to 5′ orientation. Strains CC-503 and CC-4532 assembled from PacBio reads (Craig et al., *in prep*), CC-1690 from Nanopore (O'Donnell et al. 2020).

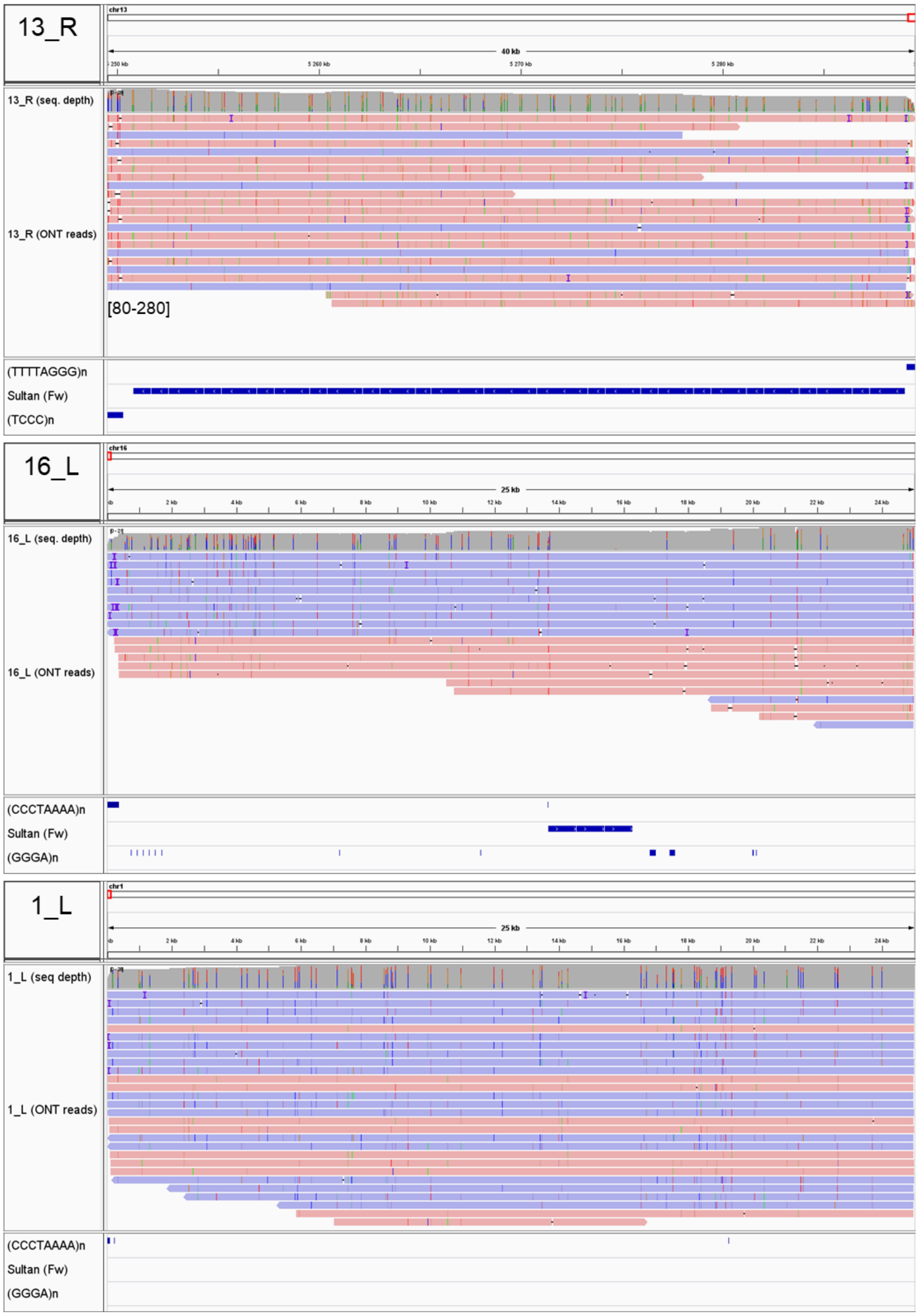


**Supplemental Figure S2. Mapping of Nanopore reads against final assembly of representative class A, C and D subtelomeres (13_R, 16_L and 1_L, respectively).**

The read dataset was down-sampled to 40x depth before alignment. The mapping used minimap2 and was visualized with IGV (http://igv.org/). For each panel, upper frame: genomic coordinates; grey chart:

read depth (vertical bars for positions with SNVs: N, A, T, G and C, in grey, green, red, orange and blue, respectively); pink and mauve horizontal arrows: Nanopore reads from the plus or minus DNA strand, respectively; vertical violet bars and horizontal black bars on reads: insertions and deletions > 10 bp, respectively; non-unique base mismatch (“quick consensus mode”) shaded by quality; the length of over-represented INDELs is indicated in brackets for 13_R; lower frame: chromosome features are depicted in blue rectangles.

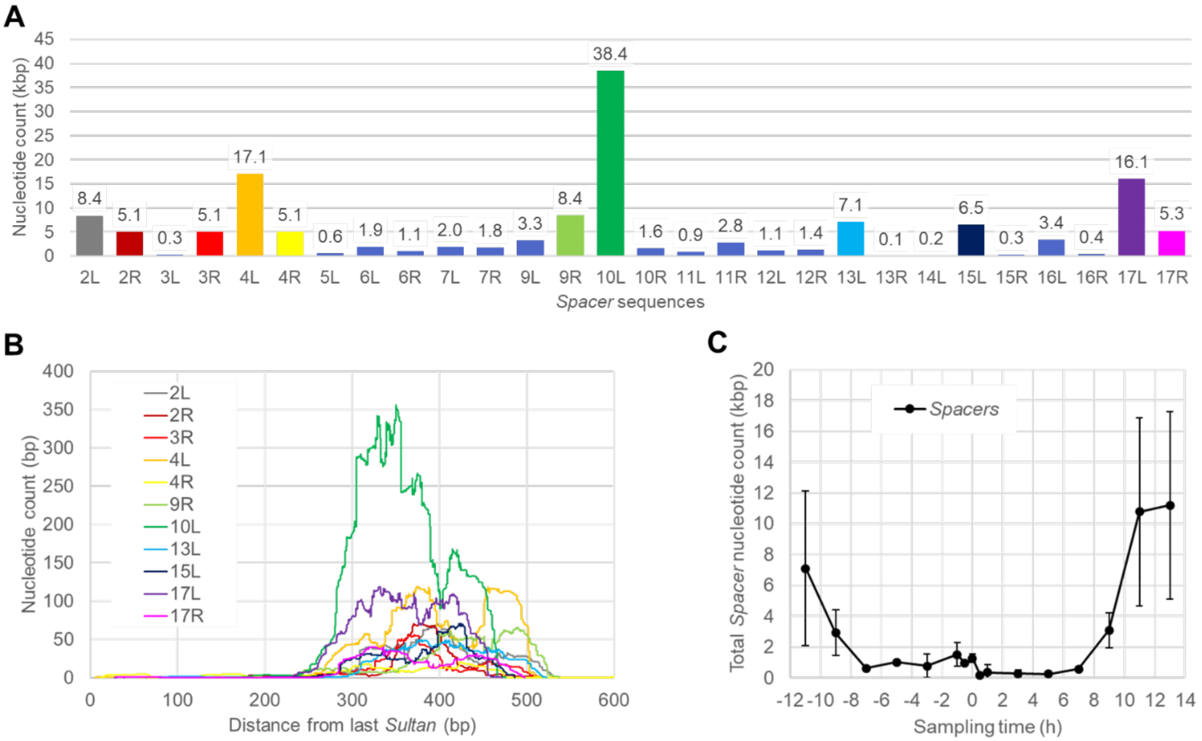


**Supplemental Figure S3. Transcription among *Spacer* sequences.**

Transcript dataset from (Strenkert et al. 2019)(accession number: GSE112394, strain CC-5390) was searched using each *Spacer* sequences as BLAST queries. *(A)* Sum of nucleotide coverage in the *Spacer* sequences across subtelomeres. *(B)* Plot of nucleotide coverage along the 11 most expressed *Spacers*, starting from the *Sultan*/*Spacer* junction (0) towards the G-rich repeats. *(C)* Mean ± SD (n = 3 biological replicates) of *Spacer* sequences expression across a diurnal cycle (day: 0-12h, see (Strenkert et al. 2019) for details).

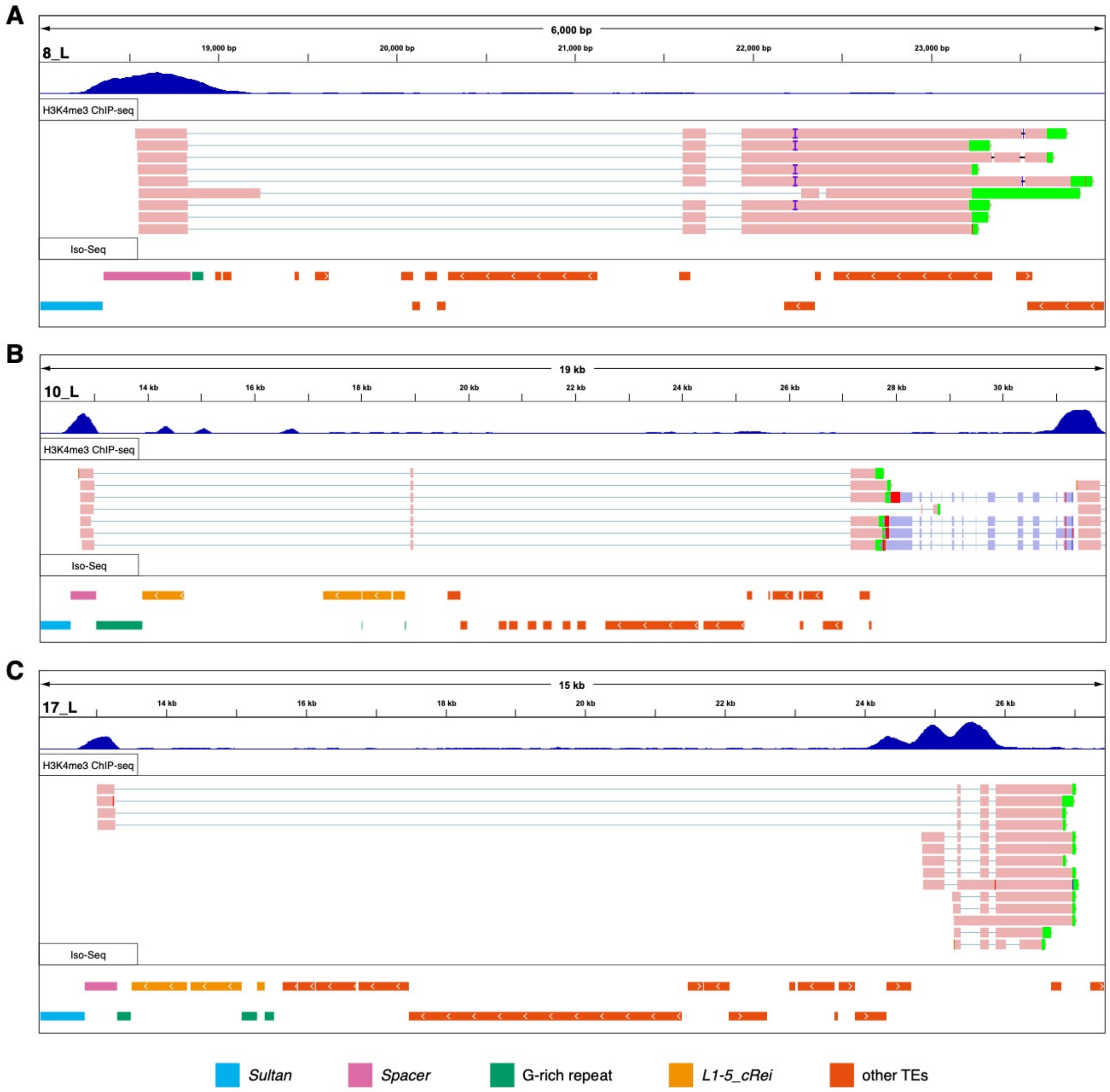


**Supplemental Figure S4. Example browser views of transcripts originated from *Spacer* sequences.**

H3K4me3 ChIP-seq (accession number: PRJNA681680) and Iso-Seq (accession number: PRJNA670202) data (Gallaher et al. 2021) were mapped to the CC-1690 assembly. Each panel shows, from top to bottom: genomic coordinates, H3K4me3 signal, individual reads and subtelomere elements (colored boxes). *(A)* Subtelomere 8_L, the final exon may be partially derived from TEs. *(B)* Subtelomere 10_L. *(C)* Subtelomere

17_L, an example of the *Spacer* forming an alternative promoter or transcription start site for a downstream gene.

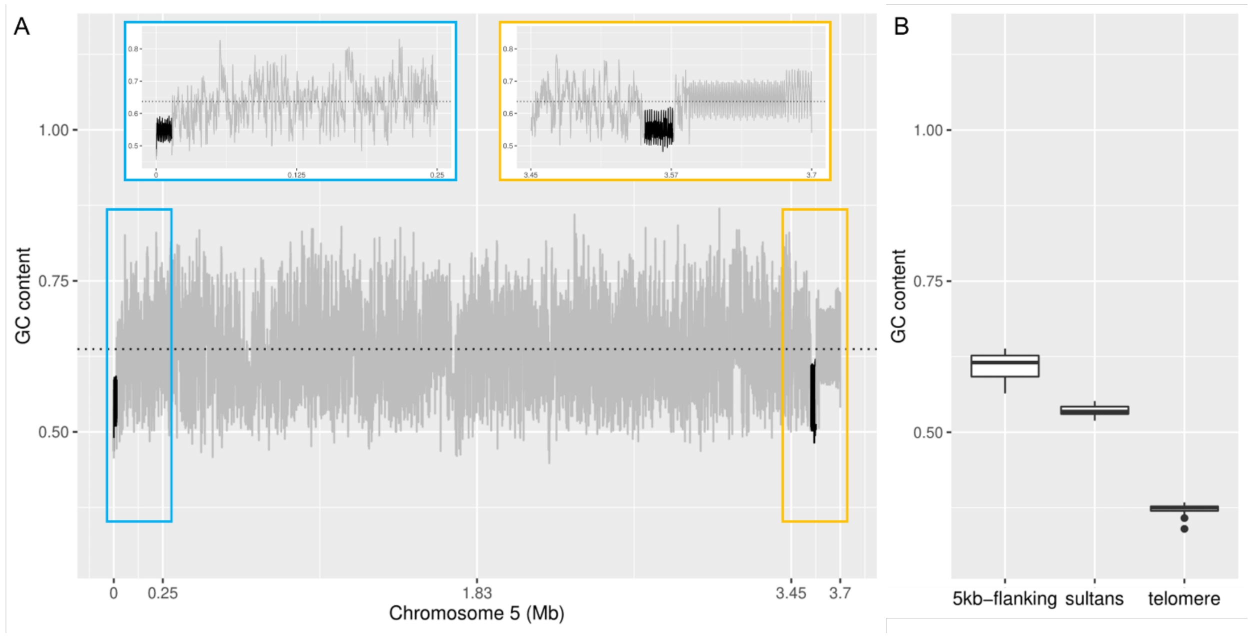


**Supplemental Figure S5. GC content in subtelomeres.**

*(A)* Example of GC content in chromosome 5 with *Sultan* repeats highlighted (dark grey). The horizontal dotted line indicates the average GC content within chromosome 5. Boxes in blue and orange highlight a 250 kb window at the chromosome extremities. Note that the 5_R contains *Suber* and *Subtile* repeats on its distal part. *(B)* Boxplot of GC content within all telomeres, *Sultan* arrays and the 5 kb surrounding *Sultan* (excluding telomere). GC content was calculated using a 500 bp window, sliding 100 bp for each window calculated.

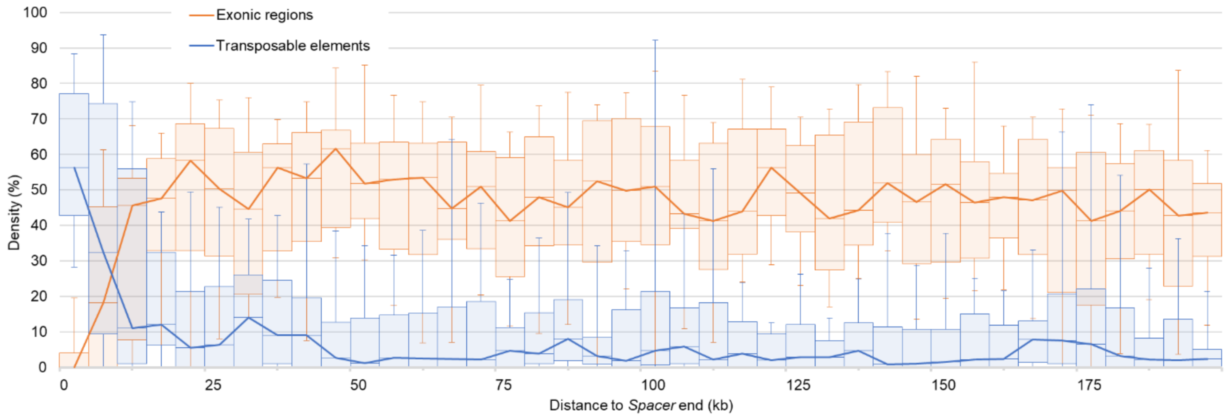


**Supplemental Figure S6. Density of exons and TEs in subtelomeres of strain CC-4532.**

Proportions of exonic regions and TEs were calculated in 5 kb windows over 200 kb (starting from the G-rich regions, “0”) for class A and class B subtelomeres. Shown are the median (line), 1st and 3rd quartile (boxes) and 1st and 9th decile (whiskers) across subtelomeres.

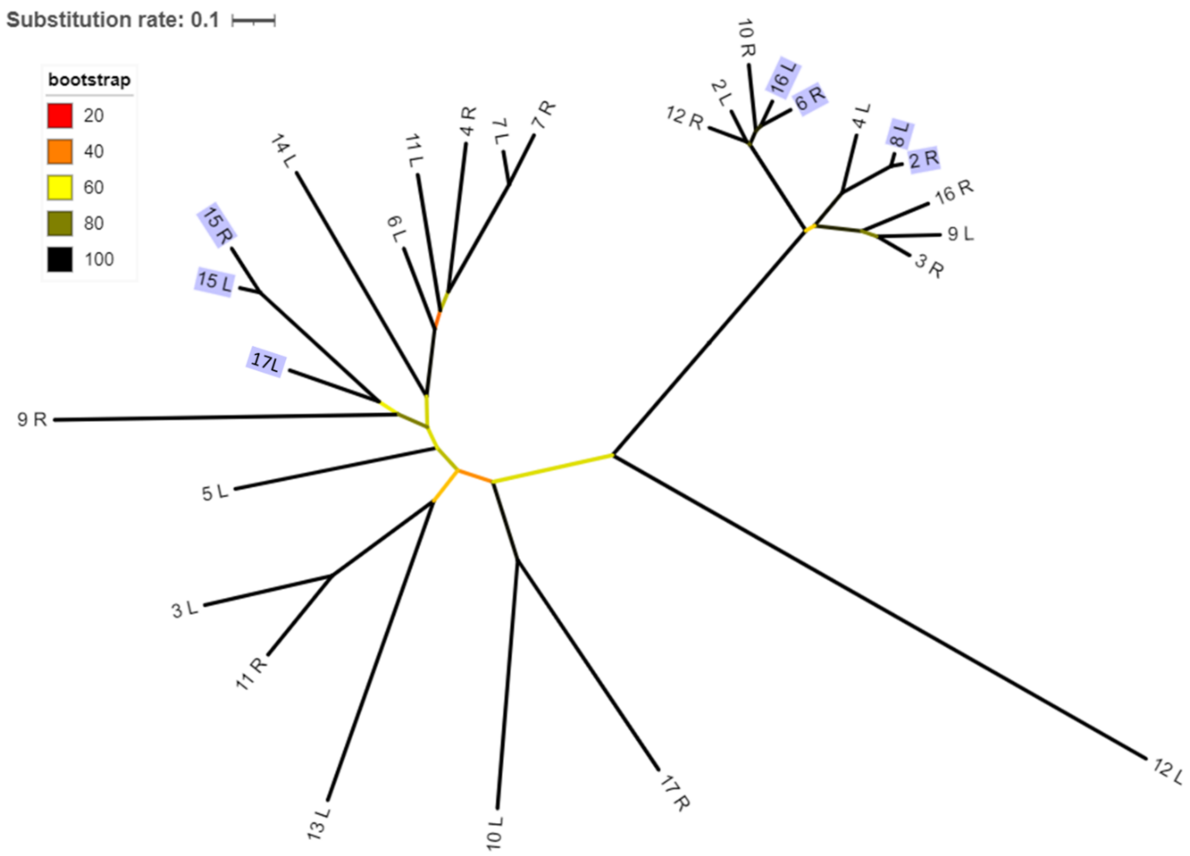


**Supplemental Figure S7. Phylogenetic tree of *Spacer* sequences from strain CC-1690.**

*Spacer* sequences were aligned using MAFFT and maximum-likelihood trees constructed with PhyML. Branches are colored from red to black according to bootstrap values. Highlighted *Spacers* are consistent with the *Sultan* repeats phylogeny.

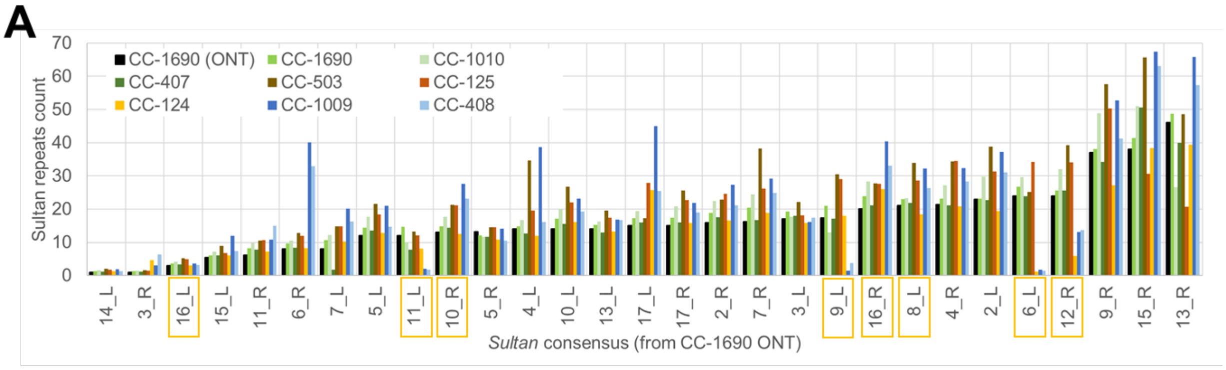


**B**

| $r^2$ | CC-1690 | CC-1010 | CC-407 | CC-503 | CC-125 | CC-124 | CC-1009 | CC-408 |
|---|---|---|---|---|---|---|---|---|
| **CC-1690** | 0,99 | 0,76 | 0,92 | 0,83 | 0,58 | 0,64 | 0,51 | 0,59 |

**Supplemental Figure S8. Count of *Sultan* repeats for each subtelomere in laboratory strains.**

*(A)* Estimate of repeat count as in Fig. 6. Subtelomeres potentially affected by the distribution of haplotype blocks among these strains are highlighted. *(B)* Determination coefficients ($r^2$) of estimated *Sultan* repeat count in laboratory strains against CC-1690 Nanopore assembly.

**Alignments.zip**

**Supplemental File F1. Alignment of all individual copies of subtelomeric repeats.**

Each individual sequences and coordinates were extracted using blast algorithm and phased. Sequences were aligned using MAFFT.

**Subtelomere_repeats_detection.sh**

**Supplemental File F2. Script Subtelomere_repeats_detection.sh.**

Script used to collect individual copies of repeats at subtelomeres.

**Illumina_Sultan_coverage.sh**

**Supplemental File F3. Script Illumina_Sultan_coverage.sh.**

Script used to map Illumina reads to reference genome and calculate coverage of subtelomere-specific *Sultan* consensus.
