## Supplemental File F2 for "Architecture and evolution of subtelomeres in the unicellular green alga *Chlamydomonas reinhardtii*": Subtelomere_repeats_detection.sh

```
##!/bin/bash -x
#usage bash /data/ovallon/scripts/TandemRepeatFinder.sh <genome name> <absolute path to genome fasta>
#check line 45 whether you want to run TRF on all sequences, or just the chromosomes (e.g. chr*_R.fa chr*_L.fa)
module load EMBOSS/6.4.0
module load TRF/4.04
module load bedtools/2.25.0
module load  blast+/2.6.0
module load mafft/7.130
genome=$1
genomepath=$2
cd /data/ovallon/Telomeres/
mkdir $genome
cd $genome
echo 'now splitting ' $genomepath
#Split fasta, making sure each entry name is a single word
awk '{print $1}' $genomepath | awk '{ if (substr($0, 1, 1)==">") {filename=(substr($0,2) ".fa")}        print $0 > filename }'
#Read molecule length:
cat *.fa |awk  'BEGIN {; OFS = "\t"} /^>/ && r { print r, s; r=s="" } /^>/ { r = substr($0, 2); next } { s += length } END { print r "\t" s }' | egrep -v 'ChrM|ChrC' > in.lengths
#Count number of molecules:
wc -l  in.lengths > in.count
#For each, extract the Left 30 kb (or less) with seqret. For this, generate a script with awk:
echo 'now extracting L and R 30 kb'
awk '{ print "seqret -sequence " $1 ".fa -sbegin1 1 -send1 130000 -outseq Molecule_ends/" $1 "_L.fa -sid1 " $1 "_L" }' in.lengths > extract_L.sh
#Extract and revcomp the right ends if molecule is longer than 130 kb (otherwise redundent with left)
awk '{ if ($2>130000) print "seqret -sequence " $1 ".fa -sbegin1 " $2-130000+1 " -send1 " $2 " -outseq Molecule_ends/" $1 "_R.fa -sreverse1 -sid1 " $1 "_R" }' in.lengths > extract_R.sh
mkdir Molecule_ends
bash extract_L.sh 
bash extract_R.sh
wait
cd Molecule_ends
cat *_R.fa *_L.fa > ALL_ends.fa
#run DUST to identify low complexity regions (ex G-rich repeats, not seen by TRF when the parameters are set to avoid duplication of the Sultan elements
#run TandemRepeatFinder on all _R.fa and _L.fa files in the directory
echo 'running Tandem Repeat Finder with stringent parameters'
for file in chr*_R.fa chr*_L.fa
do
basename=${file%%.*}
trf  ${file} 3 5 5 80 20 100 2000 &
dustmasker -in  ${file}  -linker 1 -outfmt interval -level 35  -window 64 -out ${basename}.dust &
wait
done
wait
mkdir TandemRepeatFinder
mv *html *.dust TandemRepeatFinder/
cd TandemRepeatFinder/
#extract fasta of all TRF consensuses so that each sequence reads name@start--end;period
echo 'parsing TRF output to generate fasta'
for f in *.txt.html
do
basename=${f%%.*} 
egrep 'Sequence: |Indices|Period|^[ACGT]+$' ${f} | sed -E 's/    Indices: ([0-9]*)--([0-9]*)  Score: ([0-9]*)/;\1;\2;\3/g' | sed -E 's/    Period size: ([0-9]*)  Copynumber: ([0-9\.]*)  Consensus size: ([0-9]*)/;\1;\2;\3/g' | awk '/^;/ {printf("\n%s\n",$0);next; } { printf("%s",$0);}  END {printf("\n");}' |sed  '/^$/d' | awk 'BEGIN { RS = "Sequence: " ; FS = "\n" ; OFS = "\n" } {for(i=2;i<NF;i++) if (i % 3 == 2) printf ">" $1  $i $i+1 ; else print $i }' > ${basename}_TRF_consensus.fasta
done
#extract fasta of all DUST masked sequences so that each sequence reads name@start--end;period
echo 'parsing DUST output to generate fasta'
for f in *.dust
do
basename=${f%%.*}
egrep -v '>' ${f} | awk -v a="$basename" '{print "seqret -sequence ../" a ".fa -sbegin1 " $1 " -send1 " $3 " -outseq " a "_" $1 "_" $3 "_dust_multiline.fa  -sid1 " a "_" $1 "_" $3 "_dust"}' |sed '/^$/d' >> extract_DUST.sh
done
wait
bash  extract_DUST.sh
mkdir individual_results
mv *.html *consensus.fasta *_dust_multiline.fa *.dust individual_results/
cat individual_results/*_dust_multiline.fa > ALL_ends_DUST_multiline.fasta
cat individual_results/*_TRF_consensus.fasta > ALL_ends_TRF_consensus.fasta
#convert ALL_ends_DUST.fasta to uniline
awk '/^>/ {printf("\n%s\n",$0);next; } { printf("%s",$0);}  END {printf("\n");}' < ALL_ends_DUST_multiline.fasta > ALL_ends_DUST.fasta
#identify molecules with a telomere-like repeat less than 100 nt long and starting within first 9 nt
egrep 'AAACCCT|AACCCTA|ACCCTAA|CCCTAAA|CCTAAA|CTAAA[AC]|AAAACCC|TAAACC' -B 1 ALL_ends_TRF_consensus.fasta | egrep  -A 1 '^[^;]+;.;[^;]+;[^;]+;[0-9]{1,2};' | egrep -v '^--'  > Telo_repeats.fasta
egrep '>' Telo_repeats.fasta | cut -d ';' -f 1 | sed 's/>//g' | sort -u > Telo_scaffolds.list
#identify scaffolds with a TRF but without a Telo
egrep '>' ALL_ends_TRF_consensus.fasta | cut -d ';' -f 1 | sort -u | sed 's/>//g' | egrep -v -f Telo_scaffolds.list | egrep -v '^--' > NonTelo_with_repeats.list
#identify G-rich repeats
awk '{ if ($0 ~ /^>/) print $0; else {if ( gsub(/G/, "G")> gsub(/A/, "A")+gsub(/T/, "T")+gsub(/C/, "C")) {print "G-rich",$1} else print $0} }' ALL_ends_DUST.fasta | egrep -B 1 'G-rich' | egrep -v '^--' | sed 's/G-rich //g' > G-rich_DUST.fasta
awk '{ if ($0 ~ /^>/) print $0; else {if ( gsub(/G/, "G")> gsub(/A/, "A")+gsub(/T/, "T")+gsub(/C/, "C")) {print "G-rich",$1} else print $0} }' ALL_ends_TRF_consensus.fasta | egrep -B 1 'G-rich' | egrep -v '^--' | sed 's/G-rich //g' > G-rich_TRF.fasta
##extract TRF repeats starting within first kb
#egrep '^[^;]+;[0-9]{1,3};' -A 1 ALL_ends_TRF_consensus.fasta > Repeats_before_1000.fasta
#select TRF consensuses longer than 60 nt (to exclude SDR)
awk '!/^>/ { next } { getline seq } length(seq) > 60 { print $0 "\n" seq }' ALL_ends_TRF_consensus.fasta > ALL_ends_TRF_consensus+60.fasta
#blast all long consensuses to extremities
blastn -query  ALL_ends_TRF_consensus+60.fasta  -evalue 10e-20 -subject ../ALL_ends.fa -outfmt "7 qacc qstart qend qlen sacc sstart send slen evalue bitscore length mismatch gapopen " -out ALL_ends_TRF_consensus+60-ALL_ends.blastn
#blast all long consensuses to genome
blastn -query  ALL_ends_TRF_consensus+60.fasta  -evalue 10e-20 -subject $genomepath -outfmt "7 qacc qstart qend qlen sacc sstart send slen evalue bitscore length mismatch gapopen " -out ALL_ends_TRF_consensus+60-genome.blastn
egrep -v '^#' ALL_ends_TRF_consensus+60-genome.blastn | awk '{print $0,(($7<$6)?"-":"+"), (($8-$7<$6)?$8-$7:$6), 100*$6/$8, (100*($8-$7)/$8)}' | sed 's/ /\t/g' | sort -n -k1b,1 -k5b,5 -k16,16 > ALL_ends_TRF_consensus+60-genome.blastn2
#blast all long consensuses to each other, remove selfhits and extract results for Telo scaffolds
blastn -query  ALL_ends_TRF_consensus+60.fasta  -evalue 10e-20 -subject ALL_ends_TRF_consensus+60.fasta -outfmt "7 qacc qstart qend qlen sacc sstart send slen evalue bitscore length mismatch gapopen " -out  ALL_ends_TRF_consensus+60-self.blastnraw
awk 'BEGIN {RS="BLASTN"; FS="\n"; OFS="\n"}{if ($5 ~ "# 1 hits found" || $4 ~ "# 0 hits found") next; else {$1=$3=$4=""; print $0}}' ALL_ends_TRF_consensus+60-self.blastnraw | sed '/^$/d' | awk -F '\t' '{if (($1~/#/) || ($5!=$1 || $6!=$2 || $7!=$3)) print $0}' > ALL_ends_TRF_consensus+60-self.blastn
blastn -query  ALL_ends_TRF_consensus+60.fasta  -evalue 10e-20 -subject ALL_ends_TRF_consensus+60.fasta -outfmt 0 -out  ALL_ends_TRF_consensus+60-self.aln-blastn 
egrep -v '#' ALL_ends_TRF_consensus+60-self.blastn | egrep -f Telo_scaffolds.list | awk ' $1 != $5 || $2 != $6 || $3 != $7' | sort -u > Telo_TRF-ALL_TRF.blastn
#identify hits in reverse orientation, for all TRFs and for those of Telo scaffolds only 
awk '{ if ($6 > $7) print $0 }' ALL_ends_TRF_consensus+60-self.blastn > ALL_ends_TRF_consensus+60-self-minus.blastn
awk '{ if ($6 > $7) print $0 }' Telo_TRF-ALL_TRF.blastn  >  Telo_TRF-ALL_TRF-minus.blastn
#create RevComp of Telo_TRF sequences hit in minus, so that they can be aligned to the others; add _RC to their name and make them uniline
egrep -v '#' ALL_ends_TRF_consensus+60-self-minus.blastn | cut -f 5 | awk '$1!~/^$/ {print "ALL_ends_TRF_consensus.fasta:" $1}' | sort -u > To_RevComp.seqretlist
seqret -sequence @To_RevComp.seqretlist -sreverse1  -outseq RevComped.fasta
sed -E 's/>(\S+)/>\1_RC/g' RevComped.fasta | awk '/^>/ {printf("\n%s\n",$0);next; } { printf("%s",$0);}  END {printf("\n");}' > RevComped_uniline.fasta
#mask TRF repeat regions in fasta of extremity and blast masked *.fa to the long TRF consensus to identified regions homologous to the repeats that have been missed by TRF; extract individual spacer sequences between the first 5 repeats, if longer than 4 nt
rm -f *_spacers.sh
for f in individual_results/*L_TRF_consensus.fasta individual_results/*R_TRF_consensus.fasta
do
basename=$(basename $f _TRF_consensus.fasta)
egrep '>' ${f} | sed 's/>//g' | awk -F ';' 'BEGIN {OFS="\t"} {print $1, $2, $3}' | sort -u > individual_results/${basename}.mask
bedtools maskfasta -fi ../${basename}.fa -bed individual_results/${basename}.mask -fo ../${basename}_masked.fa
blastn -query ../${basename}_masked.fa -evalue 10e-20 -subject ALL_ends_TRF_consensus+60.fasta -outfmt "7 qacc qstart qend qlen sacc sstart send slen evalue bitscore length mismatch gapopen " -out individual_results/${basename}_masked-TRFs.blastn
tr '\n' ';' < ${f} | awk -F ';' ' $2>5 {print "seqret -sequence ../" $1 "_masked.fa -sbegin1 1 -send1 " $2-1 " -outseq " $1 "_1-" $2-1 "_before_TRF_spacer.fa  -sid1 " $1 "-1-" $2-1 "-before_TRF-NA-NA-" $2-1 "nt"}' |sed 's/>//g' >> extract_before_TRF.sh
tr '\n' ';' < ${f} | awk -F ';' ' $10-$3>5 {print "seqret -sequence ../" $1 "_masked.fa -sbegin1 " $3+1 " -send1 " $10-1 " -outseq " $1 "_" $3+1 "-" $10-1 "_first_spacer.fa  -sid1 " $1 "-" $3+1 "-" $10-1 "-first_spacer-NA-NA-" $10-$3-1 "nt"}' |sed 's/>//g' >> extract_first_spacers.sh
tr '\n' ';' < ${f} | awk -F ';' ' $18-$11>5 {print "seqret -sequence ../" $1 "_masked.fa -sbegin1 " $11+1 " -send1 " $18-1 " -outseq " $1 "_" $11+1 "-" $18-1 "_second_spacer.fa -sid1 " $1 "-" $11+1 "-" $18-1 "-second_spacer-NA-NA-" $18-$11-1 "nt"}' |sed 's/>//g' >> extract_second_spacers.sh
tr '\n' ';' < ${f} | awk -F ';' ' $26-$19>5 {print "seqret -sequence ../" $1 "_masked.fa -sbegin1 " $19+1 " -send1 " $26-1 " -outseq " $1 "_" $19+1 "-" $26-1 "_third_spacer.fa -sid1 " $1 "-" $19+1 "-" $26-1 "-third_spacer-NA-NA-" $26-$19-1 "nt"}' |sed 's/>//g' >> extract_third_spacers.sh
tr '\n' ';' < ${f} | awk -F ';' ' $34-$27>5 {print "seqret -sequence ../" $1 "_masked.fa -sbegin1 " $27+1 " -send1 " $34-1 " -outseq " $1 "_" $27+1 "-" $34-1 "_fourth_spacer.fa -sid1 " $1 "-" $27+1 "-" $34-1 "-fourth_spacer-NA-NA-" $34-$27-1 "nt"}' |sed 's/>//g' >> extract_fourth_spacers.sh
tr '\n' ';' < ${f} | awk -F ';' ' $42-$35>5 {print "seqret -sequence ../" $1 "_masked.fa -sbegin1 " $35+1 " -send1 " $42-1 " -outseq " $1 "_" $35+1 "-" $42-1 "_fifth_spacer.fa -sid1 " $1 "-" $35+1 "-" $42-1 "-fifth_spacer-NA-NA-" $42-$35-1 "nt"}' |sed 's/>//g' >> extract_fifth_spacers.sh
tr '\n' ';' < ${f} | awk -F ';' ' $50-$43>5 {print "seqret -sequence ../" $1 "_masked.fa -sbegin1 " $43+1 " -send1 " $50-1 " -outseq " $1 "_" $43+1 "-" $50-1 "_sixth_spacer.fa -sid1 " $1 "-" $43+1 "-" $50-1 "-sixth_spacer-NA-NA-" $50-$43-1 "nt"}' |sed 's/>//g' >> extract_sixth_spacers.sh
tr '\n' ';' < ${f} | awk -F ';' ' $58-$51>5 {print "seqret -sequence ../" $1 "_masked.fa -sbegin1 " $51+1 " -send1 " $58-1 " -outseq " $1 "_" $51+1 "-" $58-1 "_seventh_spacer.fa -sid1 " $1 "-" $51+1 "-" $58-1 "-seventh_spacer-NA-NA-" $50-$43-1 "nt"}' |sed 's/>//g' >> extract_seventh_spacers.sh
done
wait 
bash extract_before_TRF.sh
bash extract_first_spacers.sh 
bash extract_second_spacers.sh 
bash extract_third_spacers.sh 
bash extract_fourth_spacers.sh 
bash extract_fifth_spacers.sh 
bash extract_sixth_spacers.sh
bash extract_seventh_spacers.sh
wait
#the char ; is not allowed in a bash script, was replaced by "-", so now replace back to maintain correspondence with repeat names
sed -i 's/-/;/g' *spacer.fa 
#make spacer files uniline and remove spacers made entirely of Ns (i.e. included in a TRF)
for f in *_spacer.fa
do
basename=${f%%.*}
awk '/^>/ {printf("\n%s\n",$0);next; } { printf("%s",$0);}  END {printf("\n");}' ${f} > ${basename}_uniline.fa
done
mv *_spacer*.fa individual_results/
cat individual_results/*_spacer_uniline.fa | awk 'BEGIN { RS = ">" ; FS = "\n" ; OFS = "\n" } {if ($2~/^N+$/) next; else {print ">" $1,$2 }}'> ALL_spacers.fa
cat ../*_masked.fa > ../ALL_masked.fa
blastn -query ../ALL_masked.fa -evalue 10e-20 -subject ../ALL_masked.fa -outfmt "7 qacc qstart qend qlen sacc sstart send slen evalue bitscore length mismatch gapopen " -out ALL_masked-self.blastnraw
awk 'BEGIN {RS="BLASTN"; FS="\n"; OFS="\n"}{if ($5 ~ "# 1 hits found" || $4 ~ "# 0 hits found") next; else {$1=$3=$4=""; print $0}}'  ALL_masked-self.blastnraw | sed '/^$/d' | awk -F '\t' '{if (($1~/#/) || ($5!=$1 || $6!=$2 || $7!=$3)) print $0}' >  ALL_masked-self.blastn
blastn -query ../ALL_masked.fa -evalue 10e-20 -subject ../ALL_masked.fa -outfmt 0 -out ALL_masked-self.aln-blastn
#extract regions of masked fasta that hit other masked fasta
awk -F '\t' '$1!~/#/ {print $1, $2, $3}{OFS="\t"}' ALL_masked-self.blastn | sort -u -k1,1 -k2,2n -k3,3n > ALL_masked-self.bed
bedtools merge -i ALL_masked-self.bed | awk '{print "seqret -sequence ../" $1 "_masked.fa -sbegin1 " $2 " -send1 " $3 " -outseq " $1 "-" $2 "-" $3 "-shared.fasta  -sid1 " $1 "-" $2 "-" $3 "-shared-NA-NA-" $3-$2+1 "nt"}' |sort -u > extract_shared.sh
bash extract_shared.sh
mv *shared.fasta individual_results
cat individual_results/*shared.fasta > ALL_shared.fasta
blastn -query ALL_shared.fasta -evalue 10e-20 -subject ALL_shared.fasta -outfmt "7 qacc qstart qend qlen sacc sstart send slen evalue bitscore length mismatch gapopen " -out ALL_shared-self.blastnraw
awk 'BEGIN {RS="BLASTN"; FS="\n"; OFS="\n"}{if ($5 ~ "# 1 hits found" || $4 ~ "# 0 hits found") next; else {$1=$3=$4=""; print $0}}' ALL_shared-self.blastnraw | sed '/^$/d' | awk -F '\t' '{if (($1~/#/) || ($5!=$1 || $6!=$2 || $7!=$3)) print $0}' > ALL_shared-self.blastn
egrep -v '#' ALL_shared-self.blastn | cut -f 1,5 | awk 'BEGIN{OFS="\t"}{if ($2>$1) print $1 , $2; else print $2 , $1}' | sort -u > ALL_shared-self.tab
cp ALL_shared-self.tab ALL_shared-self_sorting.tab
#manually group shared.fa sequences based on transitive pairing (see below for spacers) and run mafft. This can be done by running the below commands individually after completion of the script and manual sorting
#awk 'BEGIN{OFS="\n"}{print $1,$2}' ALL_shared-self_group<n>.tab | awk '{$1!~/^$/ print "ALL_shared.fasta:" $1}' | sort -u >  ALL_shared_group<n>.seqretlist
#seqret -sequence @ALL_shared_group<n>.seqretlist -outseq ALL_shared_group<n>.fasta
#mafft --maxiterate 3 --globalpair --thread 20 --reorder ALL_shared_group<n>.fasta > ALL_shared_group<n>.mafft
#blast spacers to themselves and to TRFs (no filtering) and clean output from self- and no-hits
blastn -query ALL_spacers.fa -dust no -evalue 10e-20 -subject ALL_spacers.fa -outfmt "7 qacc qstart qend qlen sacc sstart send slen evalue bitscore length mismatch gapopen " -out ALL_spacers-spacers.blastnraw
awk 'BEGIN {RS="BLASTN"; FS="\n"; OFS="\n"}{if ($5 ~ "# 1 hits found" || $4 ~ "# 0 hits found") next; else {$1=$3=$4=""; print $0}}' ALL_spacers-spacers.blastnraw | sed '/^$/d' | awk -F '\t' '{if (($1~/#/) || ($5!=$1 || $6!=$2 || $7!=$3)) print $0}' > ALL_spacers-spacers.blastn
blastn -query ALL_spacers.fa -dust no -evalue 10e-20 -subject ALL_spacers.fa -outfmt 0 -out ALL_spacers-spacers.aln-blastn 
blastn -query ALL_spacers.fa -dust no -evalue 10e-20 -subject ALL_ends_TRF_consensus+60.fasta -outfmt "7 qacc qstart qend qlen sacc sstart send slen evalue bitscore length mismatch gapopen " -out ALL_spacers-TRF.blastn 
#blast spacers to extremities (no filtering)
blastn -query ALL_spacers.fa -dust no -evalue 10e-20 -subject ../ALL_ends.fa -outfmt "7 qacc qstart qend qlen sacc sstart send slen evalue bitscore length mismatch gapopen " -out ALL_spacers-extremities.blastn 
#blast spacers to entire genome, filering low complexity
blastn -query ALL_spacers.fa -evalue 10e-20 -subject $genomepath -outfmt "7 qacc qstart qend qlen sacc sstart send slen evalue bitscore length mismatch gapopen " -out ALL_spacers-genome.blastn 
wait
egrep -v '^#' ALL_spacers-genome.blastn | awk '{print $0,(($7<$6)?"-":"+"), (($8-$7<$6)?$8-$7:$6), 100*$6/$8, (100*($8-$7)/$8)}' | sed 's/ /\t/g' | sort -n -k1b,1 -k5b,5 -k16,16 > ALL_spacers-genome.blastn2
#extract hits of spacers to genome that are more than 30 kb from extremity
awk -F '\t' '($15>30000) {print $0}' ALL_spacers-genome.blastn2 > ALL_spacers-genome_internal.blastn
cut -f 1 ALL_spacers-genome_internal.blastn | sort -u > spacers_with_internal_hits.list
#extract from blast only (non-self) results pertaining to extremities with Telo arrays
egrep -v '#' ALL_spacers-spacers.blastn | egrep  -f Telo_scaffolds.list > ALL_spacers-spacers-Telo_only.blastn
egrep -v '#' ALL_spacers-genome.blastn2 | egrep  -f Telo_scaffolds.list  > ALL_spacers-genome-Telo_only.blastn2
#list Telo_TRF sequences hitting another TRF sequence or a spacer (in case spacers were repeats not identified as such by TRF)
egrep -v '#' ALL_spacers-TRF.blastn | egrep -f Telo_scaffolds.list > spacers-Telo_TRF.blastn
cut -f 1,5 spacers-Telo_TRF.blastn | awk 'BEGIN{OFS="\t"}{if ($2>$1) print $1 , $2; else print $2 , $1}' | sort -u > spacers-Telo_TRF.tab
cut -f 1 spacers-Telo_TRF.blastn | awk '{print "ALL_spacers.fa:" $1}' | sort -u > spacers-hitting-Telo-TRFs.seqretlist
seqret -sequence @spacers-hitting-Telo-TRFs.seqretlist -outseq spacers-hitting-Telo-TRFs.fasta
awk '/^>/ {printf("\n%s\n",$0);next; } { printf("%s",$0);}  END {printf("\n");}' spacers-hitting-Telo-TRFs.fasta >  spacers-hitting-Telo-TRFs_uniline.fasta 
cut -f 1,5 Telo_TRF-ALL_TRF.blastn | awk 'BEGIN{OFS="\t"}{if ($2>$1) print $1 , $2; else print $2 , $1}' | sort -u > Telo_TRF_pairs.tab
#identify which Telo scaffolds have a TRF consensus similar to that of another Telo scaffold
awk -F ';|\t' 'BEGIN{OFS="\n"}{print $1, $8}' Telo_TRF_pairs.tab |sort -u > Scaffolds_with_TRFs_similar_to_subTelo_TRFs.list
#double all Telo TRF sequences and their spacer and TRF hits, including those RevComped
awk 'BEGIN{OFS="\n"}{print $1,$5}' Telo_TRF-ALL_TRF.blastn | awk '$1!~/^$/ {print "ALL_ends_TRF_consensus.fasta:" $1}' | sort -u > Telo_TRF_with_hits.seqretlist
seqret -sequence @Telo_TRF_with_hits.seqretlist -outseq  Telo_TRF_with_hits.fasta
awk '/^>/ {printf("\n%s\n",$0);next; } { printf("%s",$0);}  END {printf("\n");}' Telo_TRF_with_hits.fasta > TRF_consensus_with_hits_uniline.fasta
cat TRF_consensus_with_hits_uniline.fasta RevComped_uniline.fasta | sed -e 's/^$//g' | awk 'BEGIN { RS = ">" ; FS = "\n" ; OFS = "\n" } { print ">" $1,$2,$2 }' | sed 's/^>$//g' > TRF_with_hits_doubled.fasta
cat TRF_with_hits_doubled.fasta spacers-hitting-Telo-TRFs_uniline.fasta > Homologous_TRF+spacers.fasta
cat Telo_TRF_pairs.tab spacers-Telo_TRF.tab > TRF+spacers_pairs.tab
cp TRF+spacers_pairs.tab TRF+spacers_pairs_sorting.tab
##identify which consensuses and spacers have to be aligned together based on blast hits. This manual step can be done in vim, after creating a series of TRF_group<n>.tab files. For each group of pairs sharing a sequence, in VISUAL MODE select the lines, d, :wn, P to appropriate file TRF_group<n>.tab, :wN to previous file, until only remain pairs of sequences from the same end (these don't need to be aligned if overlapping)
##then convert each TRF_group<n>.tab into a uniqued seqretlist of unique sequences which will then be used in mafft, including those sequences that must also be considered in RC orientation. This is done as a loop using commands based on the following awk and seqret
#awk 'BEGIN{OFS="\n"}{print $1,$2}' TRF_group<n>.tab | awk '{$1!~/^$/ print "Homologous_TRF+spacers.fasta:" $1}' | sort -u >  TRF_group<n>.seqretlist
#seqret -sequence @TRF_group<n>.seqretlist -outseq TRF_group<n>.fasta
#mafft --maxiterate 3 --globalpair --thread 20 --reorder Homologous_TRF+spacers.fasta > Homologous_TRF+spacers.mafft
#for f in TRF_group*.tab
#do
#basename=${f%%.*}
#awk 'BEGIN{OFS="\n"}{print $1,$2}' ${f} | awk '$1!~/^$/ {print "Homologous_TRF+spacers.fasta:" $1}' | sort -u > ${basename}.seqretlist
#seqret -sequence @${basename}.seqretlist -outseq ${basename}.fasta
#mafft --maxiterate 1000 --localpair --thread 20 --reorder  ${basename}.fasta > ${basename}.mafft 
#done
#wait
##Manual intervention to split mafft files grouping the same sequence in RC and direct orientations: in Bioedit or other viewer, separate sequences aligning in the same orientation (=> *.tab1.mafft, *.tab2.mafft), identifying those that align to themself (inverted repeat). In mafft with most sequences, remove left and right part of alignment to maximiize overlap between homologous repeats and save as {basename}_phased.mafft. 
##Generate separate fasta files for each type. Using browser, record at which position in the final fasta the first instance starts and the last instance ends and write it in defline
##select most representtative phased TRF_consensuses as Selected_phased-consensuses.fasta
##blast repeats to the genome and calculate distance to closest extremity
#blastn -query  Selected_phased-consensuses.fasta  -evalue 10e-20 -subject $genomepath -outfmt "7 qacc qstart qend qlen sacc sstart send slen evalue bitscore length mismatch gapopen " -out  Selected_phased-consensuses-genome.blastn
#egrep -v '^#' Selected_phased-consensuses-genome.blastn | awk '{print $0,(($7<$6)?"-":"+"), (($8-$7<$6)?$8-$7:$6), 100*$6/$8, (100*($8-$7)/$8)}' | sed 's/ /\t/g' | sort -n -k1b,1 -k5b,5 -k16,16 > Selected_phased-consensuses-genome.blastn2
#awk ' $7>$6 {print "seqret -sequence ../../" $5 ".fa -sbegin1 " $6 " -send1 " $7 " -outseq individual_results/" $5 "_" $6 "-" $7 "_I_copy.fasta -sid1 " $5 "_" $6 "-" $7} $6>$7 {print  "seqret -sequence ../../" $5 ".fa -sbegin1 " $7 " -send1 " $6 " -outseq individual_results/" $5 "_" $7 "-" $6 "_I_copy.fasta -sid1 " $5 "_" $7 "-" $6 "_reversed -sreverse1 "}' Selected_phased-consensuses-genome.blastn2 > retrieve_copies.sh
##Because the above command may truncate some of the copies, blast all consensuses of a given type (here Type-n) to the genome and retrieve separately the merged intervals for retrieval of all complete copies
#blastn -query TRF_group<n>_phased.fasta -evalue 10e-20 -subject /data/ovallon/Telomeres/Samuel/chlamy_nanopore_refined.fa -outfmt "7 qacc qstart qend qlen sacc sstart send slen evalue bitscore length mismatch gapopen " -out Type<n>_ALL-Nanopore_genome.blastn
#awk -F '\t' '$1!~/#/ {print $5, $6, $7}{OFS="\t"}' TypeI_ALL-Nanopore_genome.blastn | sort -u -k1,1 -k2,2n -k3,3n > TypeI_ALL-Nanopore_genome.bed
#awk '$3<$2 {print $1, $3, $2}' TypeI_ALL-Nanopore_genome.bed | sed 's/ /\t/g' | sort -u -k1,1 -k 2,2n -k 3,3n > TypeI_ALL-Nanopore_genome_minus.bed
#awk '$3>$2 {print}' TypeI_ALL-Nanopore_genome.bed > TypeI_ALL-Nanopore_genome_plus.bed
#bedtools merge -i TypeI_ALL-Nanopore_genome_minus.bed | awk '{print "seqret -sequence ../../" $1 ".fa -sbegin1 " $2 " -send1 " $3 " -outseq " $1 "-" $2 "-" $3 "_reversed_TypeI_copy.fasta  -sid1 " $1 "-" $2 "-" $3 "_TypeI_copy_reversed_" $3-$2+1 "nt -sreverse1"}' |sort -u -k3,3 -k5,5n -k 7,7n > retrieve_TypeI_copies_minus.sh
#bedtools merge -i TypeI_ALL-Nanopore_genome_plus.bed | awk '{print "seqret -sequence ../../" $1 ".fa -sbegin1 " $2 " -send1 " $3 " -outseq " $1 "-" $2 "-" $3 "_TypeI_copy.fasta  -sid1 " $1 "-" $2 "-" $3 "_TypeI_copy_" $3-$2+1 "nt"}' |sort -u -k3,3 -k5,5n -k 7,7n > retrieve_TypeI_copies_plus.sh
#bash retrieve_TypeI_copies_minus.sh &
#bash retrieve_TypeI_copies_plus.sh &
#wait
#mv *TypeI_copy.fasta individual_results/
#cat individual_results/*TypeI_copy.fasta > ALL_TypeI_copies.fasta
#mafft --maxiterate 1000 --localpair --thread 20 --reorder ALL_TypeI_copies.fasta > ALL_TypeI_copies.mafft
#
#bash retrieve_copies.sh
#cat individual_results/*_I_copy.fasta > ALL_I_copies.fasta
#mafft --maxiterate 1000 --localpair --thread 20 ALL_I_copies.fasta > ALL_I_copies.mafft
## With Bioedit, calculate distance matrix between copies using Alignment/Sequence identity matrix,  to see if copies within an array are closer to each other than across arrays.
## With MEGA-X, generate bootstrapped Maximum Likelyhood trees for the repeat consensuses and the spacers, to see if their evolution is congruent

#generate general statistics
echo "Analysis of $genome in file $genomepath" > ${genome}"_stats.txt"
egrep -c '>' *.fasta >> ${genome}"_stats.txt"
egrep -c '>' *.mafft >> ${genome}"_stats.txt"
wc -l *list >> ${genome}"_stats.txt"
wc -l *tab  >> ${genome}"_stats.txt"
wc -l *sh  >> ${genome}"_stats.txt"
wc -l *blastn  >> ${genome}"_stats.txt"
wc -l *blastn2  >> ${genome}"_stats.txt"
#for each extremity, group main results in a single file
for f in ../*L.fa ../*R.fa
do
filename=${f##*/}
basename=${filename%%.*}
echo "fullfile is ${f}"
echo "filename is ${filename}"
echo "Summary for ${basename}" >  individual_results/${basename}.summary
echo "Summary for ${basename}" >  individual_results/${basename}.summary
echo "#########################" >>  individual_results/${basename}.summary
egrep -c "${basename}" *.list| awk 'BEGIN {print "----- Counts -----"} {print $0}' >>  individual_results/${basename}.summary
egrep -c "${basename}" *.blastn* >>  individual_results/${basename}.summary
egrep -c "${basename}" *.fa* >>  individual_results/${basename}.summary
echo "#########################" >>  individual_results/${basename}.summary
egrep "${basename}" Telo_scaffolds.list | awk 'BEGIN {print "----- Lists -----"} { print "Telo_scaffolds.list:", $1}' >> individual_results/${basename}.summary
#awk -v basename='${basename}' '{if ($0 ~ /basename/) print "awk Telo_scaffolds.list:", $0}' Telo_scaffolds.list >> individual_results/${basename}.summary
echo "egreped '${basename}' from Telo_scaffolds.list"
egrep "${basename}" NonTelo_with_repeats.list | awk '{print "NonTelo_with_repeats.list:", $1}' >> individual_results/${basename}.summary
egrep "${basename}" Scaffolds_with_TRFs_similar_to_subTelo_TRFs.list | awk '{print "Scaffolds_with_TRFs_similar_to_subTelo_TRFs.list:", $1}' >> individual_results/${basename}.summary
egrep "${basename}" spacers_with_internal_hits.list | awk '{print "spacers_with_internal_hits.list:", $1}' >> individual_results/${basename}.summary
echo "#########################" >>  individual_results/${basename}.summary
echo "----- TRF consensus sequences from ${basename} --------" >>  individual_results/${basename}.summary
awk 'BEGIN { RS = ">" ; FS = "\n" ; OFS = "\n" } {if ($1~/'${basename}'/) print ">" $0 }' ALL_ends_TRF_consensus.fasta | sed '/^$/d' >> individual_results/${basename}.summary
echo "----- G-rich DUST low-complexity sequences from ${basename} --------" >>  individual_results/${basename}.summary
awk 'BEGIN { RS = ">" ; FS = "\n" ; OFS = "\n" } {if ($1~/'${basename}'/) print ">" $0 }' G-rich_DUST.fasta | sed '/^$/d' >> individual_results/${basename}.summary
echo "----- Spacer sequences from ${basename} --------" >>  individual_results/${basename}.summary
awk 'BEGIN { RS = ">" ; FS = "\n" ; OFS = "\n" } {if ($1~/'${basename}'/) print ">" $0 }' ALL_spacers.fa | sed '/^$/d' >> individual_results/${basename}.summary
echo "----- Shared sequences from ${basename} --------" >>  individual_results/${basename}.summary
awk 'BEGIN { RS = ">" ; FS = "\n" ; OFS = "\n" } {if ($1~/'${basename}'/) print ">" $0 }' ALL_shared.fasta | sed '/^$/d' >> individual_results/${basename}.summary
echo "retrieved '${basename}' from fasta files"
echo "#########################" >>  individual_results/${basename}.summary
echo "----- TRF consensus self Blastn, results for ${basename} --------" >>  individual_results/${basename}.summary
egrep  "${basename}" ALL_ends_TRF_consensus+60-self.blastn | sed '/^$/d' >> individual_results/${basename}.summary
echo "----- TRF consensus self Blastn, minus orientation, results for ${basename} --------" >>  individual_results/${basename}.summary
egrep  "${basename}" ALL_ends_TRF_consensus+60-self-minus.blastn | sed '/^$/d' >> individual_results/${basename}.summary
echo "----- TRF consensus Blastn to ALL_ends, results for ${basename} --------" >>  individual_results/${basename}.summary
egrep  "${basename}" ALL_ends_TRF_consensus+60-ALL_ends.blastn | sed '/^$/d' >> individual_results/${basename}.summary
echo "----- TRF consensus Blastn to genome, results for ${basename} --------" >>  individual_results/${basename}.summary
egrep  "${basename}" ALL_ends_TRF_consensus+60-genome.blastn2 | sed '/^$/d' >> individual_results/${basename}.summary
echo "----- Blastn of spacers to themselves, results for ${basename} --------" >>  individual_results/${basename}.summary
egrep  "${basename}" ALL_spacers-spacers.blastn >> individual_results/${basename}.summary
echo "----- Blastn of spacers to TRF consensues, results for ${basename} --------" >>  individual_results/${basename}.summary
egrep  "${basename}" ALL_spacers-TRF.blastn >> individual_results/${basename}.summary
echo "----- Blastn of spacers to extremities, results for ${basename} --------" >>  individual_results/${basename}.summary
egrep  "${basename}" ALL_spacers-extremities.blastn >> individual_results/${basename}.summary
echo "----- Blastn of spacers to genome, results for ${basename} --------" >>  individual_results/${basename}.summary
egrep  "${basename}" ALL_spacers-genome.blastn2 >> individual_results/${basename}.summary
echo "----- Blastn of TRF-masked sequence to each other, results for ${basename} --------" >>  individual_results/${basename}.summary
egrep  "${basename}" ALL_masked-self.blastn >> individual_results/${basename}.summary
echo "----- Blastn of shared sequences to each other, results for ${basename} --------" >>  individual_results/${basename}.summary
egrep  "${basename}" ALL_shared-self.blastn >> individual_results/${basename}.summary
echo "#########################" >>  individual_results/${basename}.summary
wait
done
zip ../../${genome}.zip *.list *.fa* *.blastn* *.tab *.mafft *.stats individual_results/*.summary
```
