## Supplemental File F3 for "Architecture and evolution of subtelomeres in the unicellular green alga *Chlamydomonas reinhardtii*": Illumina_Sultan_coverage.sh

```
##### In order to calculate the relative coverage of the genome for a particular region(s) of interest, here the sultans) ######

##	required tools:
##	bwa
##	samtools
##	bedtools

### modify variables for each run ###
## example of running script using CC-1690 illumina data against the CC-1690 genome
## prefix for run
strain="CC1690"
## genome for alignment of illumina
reference_genome="chlamy_nanopore_refined.fa"
## tag to keep track of genome used
suffix="refCC1690"
##paired end illumina
illumina_1="illumina_data/${strain}/*_1.fastq.gz"
illumina_2="illumina_data/${strain}/*_2.fastq.gz"
## number of threads for all tools
threads="6"

### Calculating average genome wide coverage using illumina ###
## index the genome for mapping with bwa-mem
bwa index ${reference_genome}
## mapping of illumina reads
bwa mem -v 0 -t $threads ${reference_genome} \
	$illumina_1 $illumina_2 | samtools sort -o ${strain}.illumina.sorted.bam -
## indexing reference genome for samtools
samtools faidx ${reference_genome}
## getting stats of mapping
samtools flagstat ${strain}.illumina.sorted.bam > ${strain}.flagstat.txt
## get coverage only in blocks which contain the same amount of reads (-bg) (see: https://bedtools.readthedocs.io/en/latest/content/tools/genomecov.html)
## then needs to be treated differently to calculate average coverage
bedtools genomecov -bg -ibam ${strain}.illumina.sorted.bam | awk '{print $1"\t"$2"\t"$3"\t"$4"\t"$3-$2"\t"$4*($3-$2)}' | gzip > ${strain}.COV.csv.gz
##calculate average across the whole genome and assign to a variable
average=$(zcat ${strain}.COV.csv.gz | awk '{print $5"\t"$6}' | awk '{ total += $1; totall += $2 } END { print totall/total }')

### extracting illumina reads associated with sultans ###
## index bam for samtools
samtools index -@ threads ${strain}.illumina.sorted.bam
## subset the bam regions associated with the sultan regions in CC-1690 (named here: 'CC-1690_sultans.bed')
## output of a new bam per region
cat ./CC1690_sultans.bed | sed 's/\,/\t/g' | awk '{print $1":"$2"-"$3}' | while read position
do
	count=$(( $count + 1 ))
	echo $count
	echo $position
	samtools view -@ threads -b ${strain}.illumina.sorted.bam $position > $count.TEMP.bam
done
## merge all the bam files subsetted
samtools merge -@ threads ${strain}_CC1690_sultans.bam *.TEMP.bam
## remove individual subsets
rm  *.TEMP.bam
## extract all reads assocaited with this bam subset/sultan associated into a fastq file (compressed)
samtools fastq -@ threads ${strain}_CC1690_sultans.bam | gzip > ${strain}_CC1690_sultans.fq.gz


### re-map reads against sultan consensus and calculate fold increase in coverage per consesnsus ###
## index multi-fasta with all the consensus sequences (named here 'CC1690_sultans_consensus.fa') for bwa
bwa index CC1690_sultans_consensus.fa
## map illumina reads against consensus sequences
bwa mem -v 0 -t $threads CC1690_sultans_consensus.fa \
	${strain}_CC1690_sultans.fq.gz > ${strain}_CC1690_sultan.illumina.sam
## look at mapping stats
samtools flagstat ${strain}_CC1690_sultan.illumina.sam > ${strain}_CC1690_sultan.flagstat.txt
## index sultans consensus sequences for samtools
samtools faidx CC1690_sultans_consensus.fa
## transform sam to bam while removing reads that did not map (-F 4)
samtools view -@ threads -F 4 -bt CC1690_sultans_consensus.fa.fai  \
	${strain}_CC1690_sultan.illumina.sam > ${strain}_CC1690_sultan.illumina.bam
## sort bam based on genome coordinates
samtools sort -@ threads ${strain}_CC1690_sultan.illumina.bam -o ${strain}_CC1690_sultan.illumina.sorted.bam
## remove intermediate files
rm ${strain}_CC1690_sultan.illumina.sam
rm ${strain}_CC1690_sultan.illumina.bam
## get coverage for the genome at base pair basis (-d)
bedtools genomecov -d -ibam ${strain}_CC1690_sultan.illumina.sorted.bam > ${strain}_CC1690_sultan.COV_TEMP.csv
## divide the coverage at each base pair by the genome wide average to get the fold increase
cat ${strain}_CC1690_sultan.COV_TEMP.csv | awk -v average="$average" '{print $0"\t"$3/average}' > ${strain}_CC1690_sultan.COV.csv
## remove intemediate file
rm ${strain}_CC1690_sultan.COV_TEMP.csv

### results contained in file : '${strain}_CC1690_sultan.COV.csv'
```
